# Small molecule inhibitor combination treatment effectively represses global B-cell signaling in diffuse large B-cell lymphoma

**DOI:** 10.1101/2024.11.13.623398

**Authors:** Melde Witmond, Babet Bollen, Pam Heijmans, Annemiek B. van Spriel, Jessie A.L.G. van Buggenum, Wilhelm T.S. Huck

**Affiliations:** Department of Physical-Organic Chemistry, Institute for Molecules and Materials (IMM), Radboud University Nijmegen, Nijmegen, The Netherlands; Department of Medical BioSciences, Radboud University Medical Center, Radboud Institute for Medical Innovation, Nijmegen, The Netherlands; Brightlands Maastricht Health Campus, Maastricht, The Netherlands (current working address)

## Abstract

Cells sense their environment via signaling networks and malignant cells often hijack signaling pathways for their growth, which complicates defining the effects of drugs. Here, we map the state of the signaling network of B-cell lymphoma cells via simultaneous quantification of 111 (phospho-)proteins. We demonstrate that the B-cell signaling network can be disrupted with specific clinical small molecule inhibitors (iBTK, iSYK, iNFκB), thereby inducing a repressed state of the network. Principal component analysis identifies how the three inhibitors work along subtly different repression axes through the signaling state landscape. Finally, we observe that 1 µM combination treatment with all three inhibitors is more effective in inducing the repressed state than 10 µM of the single inhibitors. These results emphasize that cellular signaling occurs in complex networks, and underscore how quantification of the signaling state landscape provides insights into how combination drug treatments can bring cells in a desired network state.

## Main

Cells process information from their environment through intracellular signaling pathways. Classically, these pathways are perceived as linear cascades: signaling activation occurs upon binding of an extracellular ligand to its receptor, which triggers a cascade of sequentially activated signaling proteins, until the signal is propagated to the effector proteins^1–3^. For instance, Nunns *et al.* (2018) have shown that three signaling pathways with distinct architectures all act as linear input-output transmitters^3^. However, high-throughput datasets have elucidated that this linear cascade paradigm is often too simplistic as direct and indirect crosstalk occurs between many signaling pathways, and that a signaling network view with various branches and modules is more appropriate^2,4–6^.

In the linear cascade representation, the B-cell receptor (BCR) signaling pathway initiates with antigen engagement of the receptor followed by activation of membrane-proximal kinases LYN, SYK, and BTK, propagating until downstream effector proteins such as NFκB, ERK or p38 are activated^7,8^. However, many signaling proteins, including SYK, BTK, and PLCγ are highly interconnected and form a multitude of feedback and feedforward loops and interact with other signaling pathways^7,9–13^ (Fig. 1A and Suppl. Fig. S1). For instance, active BCR signaling through SYK is required for optimal Toll-like receptor signaling^9^, and BTK is required for chemokine-mediated cell migration^11^.

**Figure 1:**
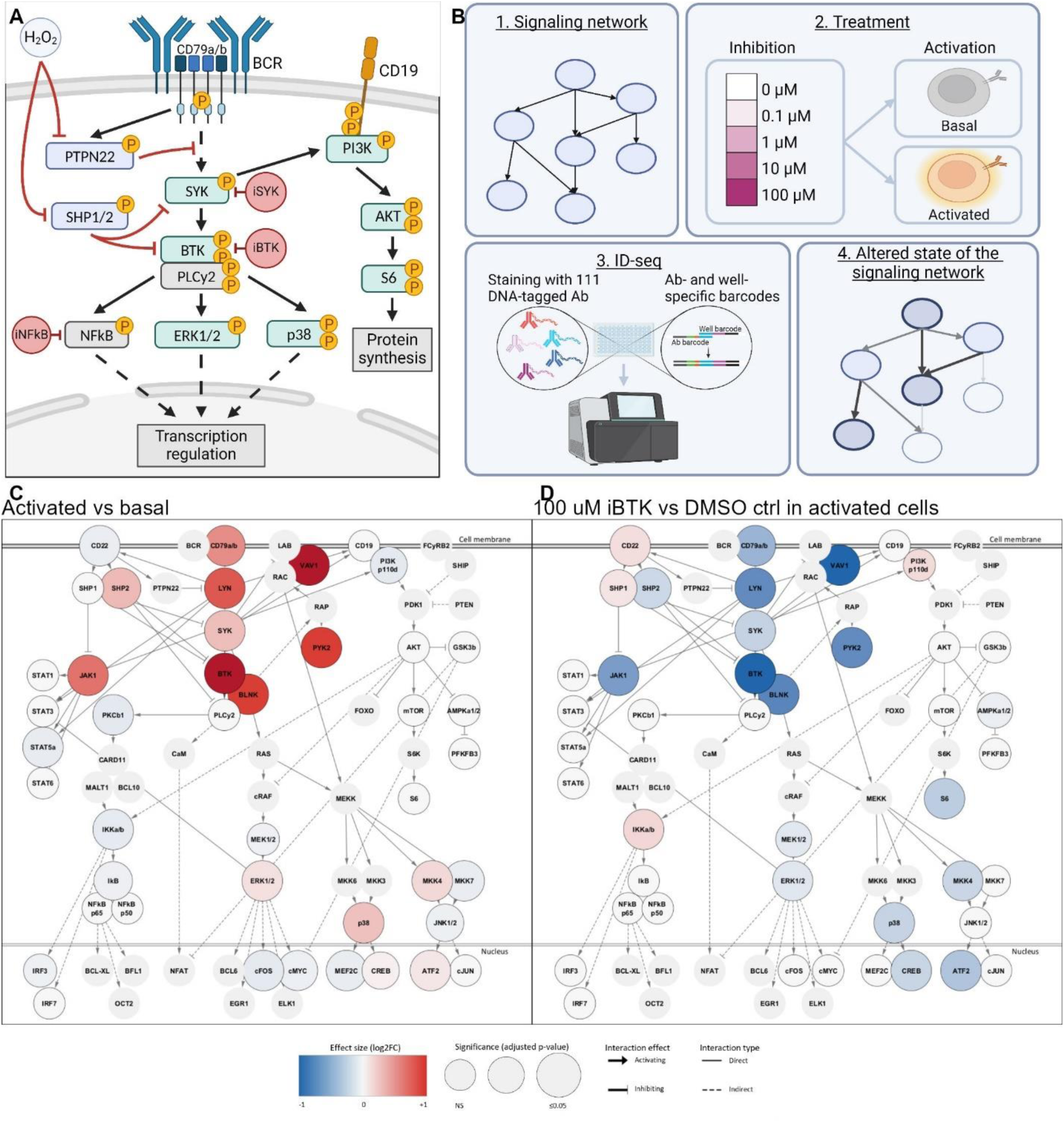
Characterization of signaling activation and inhibition by iBTK across the B-cell signaling network. A. Brief schematic overview of the B-cell signaling network, with effects of H_2_O_2_ (grey) and small molecule inhibitors indicated (red). Green = kinase, blue = phosphatase, grey = other, sharp black arrows = activation, blunt red arrows = inhibition. Adapted from Witmond et al. (2024)^45^. B. Workflow for studying signaling states. A cocktail of 15 μg/mL anti-Ig F(ab)’2 fragment, and 15 mM hydrogen peroxide (H_2_O_2_) was used to induce an activated cell state, PBS was used for the basal state. C. Network visualization of differential expression analysis (DEA) of activated cells compared to resting cells. N = 3 biological replicates per condition. Wald test with Benjamini-Hochberg (BH)-adjusted p-value threshold ≤ 0.05. D. Network visualization of DEA of 100 µM iBTK treatment compared to DMSO control in activated cells. N = 3 biological replicates per condition. Wald test with BH-adjusted p-value threshold ≤ 0.05. See Suppl. Fig. 1 for a larger schematic of the network in Fig. 1C+D.

As signaling is often organized in complex, interconnected networks, it has been proposed as early as 1999 that we should study the state of the overall network rather than individual signaling proteins - nodes in the signaling network - to understand how cells respond to various input conditions and perturbations^14,15^. As malignant cells rely on intracellular signaling pathways for their growth and survival, a network approach to understand signaling may also aid in the development and evaluation of clinical drugs, since the inhibition of a single node in an interconnected network can disrupt the balance across a large part of a signaling pathway. The seminal study of Rukhlenko *et al.* (2022) demonstrated purposeful control over cell state transitions by first mapping various cell states, modeling transitions between these states and then predicting perturbations that push cells from one state into another^16^. Similarly, to unravel the mode of action of inhibitory drugs on signaling, quantification of the complete activity state of the signaling network (that is the specific levels of protein phosphorylation across the network) is crucial. As current methods do not map changes in the overall signaling state, many disease models and drugs lack a clear view of how drugs affect cellular signaling.

Here, we studied the interconnectivity of the B-cell receptor (BCR) signaling network in context of diffuse large B-cell lymphoma (DLBCL) and treatment with three small molecule inhibitory drugs. DLBCL is a common and aggressive type of mature B-cell lymphoma^17,18^. Up to 40% of DLBCL patients do not respond or relapse after first line treatment with chemotherapy and immunotherapy. In particular, the activated B-cell (ABC) subtype of DLBCL is dependent on dysfunctional BCR and NFκB signaling for survival^8,10,19^. To obtain a complete view of the signaling state of the BCR network and examine the effects of drugs on this signaling state, we employed immunodetection by sequencing (ID-seq), a method that enables simultaneous quantification of hundreds of (phospho-)proteins^20^. In our study, we investigated three therapeutic inhibitors targeting signaling proteins BTK, SYK and NFκB. BTK can be effectively blocked by ibrutinib (referred to hereafter as iBTK)^21,22^, an inhibitor that is clinically approved for several hematological malignancies^23–27^ and under investigation for the treatment of DLBCL^28–31^. The SYK inhibitor R406 (precursor: fostamatinib; iSYK) is approved for immune thrombocytopenia treatment^32^ and under clinical investigation for DLBCL^33–35^. Lastly, the inhibitor QNZ (EVP4593; iNFκB) is a pre-clinical inhibitor described to block NFκB transcriptional activity^36,37^, which is a relevant target in ABC-DLBCL due to its dependency on NFκB signaling. Exactly how these three therapeutic drugs affect the B-cell signaling state in context of DLBCL is unknown.

We performed comprehensive ID-seq experiments with varying inhibitor doses and combinations, which allowed us to define a DLBCL-specific signaling state landscape. Our experiments elucidate that the three inhibitors work along subtly different repression axes through the signaling state landscape. We demonstrated that combination treatment with low doses of iBTK, iSYK and iNFκB together propels the B-cell network towards a repressed signaling state more effectively than high doses of the single inhibitors. These findings improve our understanding of cell signaling organization into complex and interconnected networks and highlight the ability of (combination) cancer drug treatment to induce desired signaling network states.

## Results

### ID-seq enables detailed characterization of B-cell activation and inhibition by iBTK

To determine the signaling state and interconnectivity of the B-cell signaling network in different cellular contexts, we selected the HBL1 cell line as ABC-DLBCL model and treated the cells with BTK inhibitor ibrutinib at varying doses (0.1-100 µM) in the context of basal and activated signaling (Fig. 1B). To activate B-cell signaling, cells were treated with a combination of anti-Ig F(ab)’2 fragment, to activate the BCR receptor, and hydrogen peroxide (H_2_O_2_) to inhibit phosphatases. Anti-Ig is classically used as proxy for antigen-BCR engagement^38–40^; H_2_O_2_ is a crucial second messenger in BCR signaling and present in the lymphoma tumor microenvironment^41–44^. Phospho flow cytometry confirmed that the activation with both anti-Ig and H_2_O_2_ induced significantly higher phosphorylation levels of CD79a and SYK than anti-Ig or H_2_O_2_ alone (Suppl. Fig. S2).

We used immunodetection by sequencing (ID-seq), a method that combines antibody-based target recognition and DNA sequencing through DNA-tagged antibodies^20^ (Suppl. Fig. S3), to measure the phosphorylation state of the B-cell network upon activating and inhibiting treatments. For this, we composed a panel of 111 antibodies that target total protein and phospho-protein levels of B-cell specific signaling proteins, downstream signaling pathways (NFκB, MAPK, PI3K, JAK/STAT), cell cycle proteins, and apoptosis markers (Suppl. Fig. S1 and Suppl. Table S1 for targets and detailed antibody information). The measured (phospho-)proteins in ID-seq included B-cell surface markers and components of the BCR signaling network such as CD79a (receptor complex), LYN and BTK (core BCR signaling hub), SHP2 (phosphatase), VAV1 (BCR internalization), and S6 (protein synthesis) (Fig. 1A and Suppl. Fig. S1). To confirm the ID-seq findings, we demonstrated phosphorylation levels of CD79a using phospho flow cytometry (Suppl. Fig. S4).

Next, we investigated the network-wide signaling response upon activation and inhibition with 100 µM iBTK. Differential (phospho-)protein expression analysis (DEA) provides a network-wide view of the signaling state which was used to compare the basal (resting) and activated conditions (Fig. 1C). This analysis demonstrated significantly increased phosphorylation levels of the core BCR signaling hub, consisting of CD79a, LYN, SYK, BTK, and BLNK, as well as increased phospho-levels of VAV1 and PYK2, which are involved in BCR internalization. In addition, downstream effector proteins ERK1/2, p38, and MKK4 were increased to a lesser extent. Treatment with 100 µM iBTK in the activated state resulted in substantially decreased phosphorylation of the core BCR proteins (Fig. 1D). Notably, CD22, an inhibitory receptor, showed significantly decreased phosphorylation upon activation, and iBTK treatment reversed the response to a significant increase. DMSO and PBS treatment resulted in no significant changes in signaling protein levels, indicating that changes in signaling proteins were due to inhibition by iBTK (Suppl. Figure S5). Thus, we found that phosphorylation levels of proteins across the entire B-cell network were disrupted by iBTK treatment.

### B-cell activation and inhibition by iBTK results in distinct signaling states

To obtain a generalized and dimensionality-reduced representation of the signaling states, we employed principal component analysis (PCA). The signaling state landscape was first mapped with all input conditions (Suppl. Fig. S6), and conditions of interest were subsequently sampled for visualization. PCA confirmed that BCR activation induced a highly activated signaling state that is distinct from the basal state of the B-cell signaling network (Fig. 2A, captured mostly in PC1; see Suppl. Fig. S7 for PC1 to PC3). Moreover, treatment with 100 µM iBTK pushed the signaling network towards a repressed state: signaling did not revert back to the basal state upon iBTK treatment but instead reached a new, repressed state, even for the basal-inhibited condition (captured mostly in PC3).

**Figure 2:**
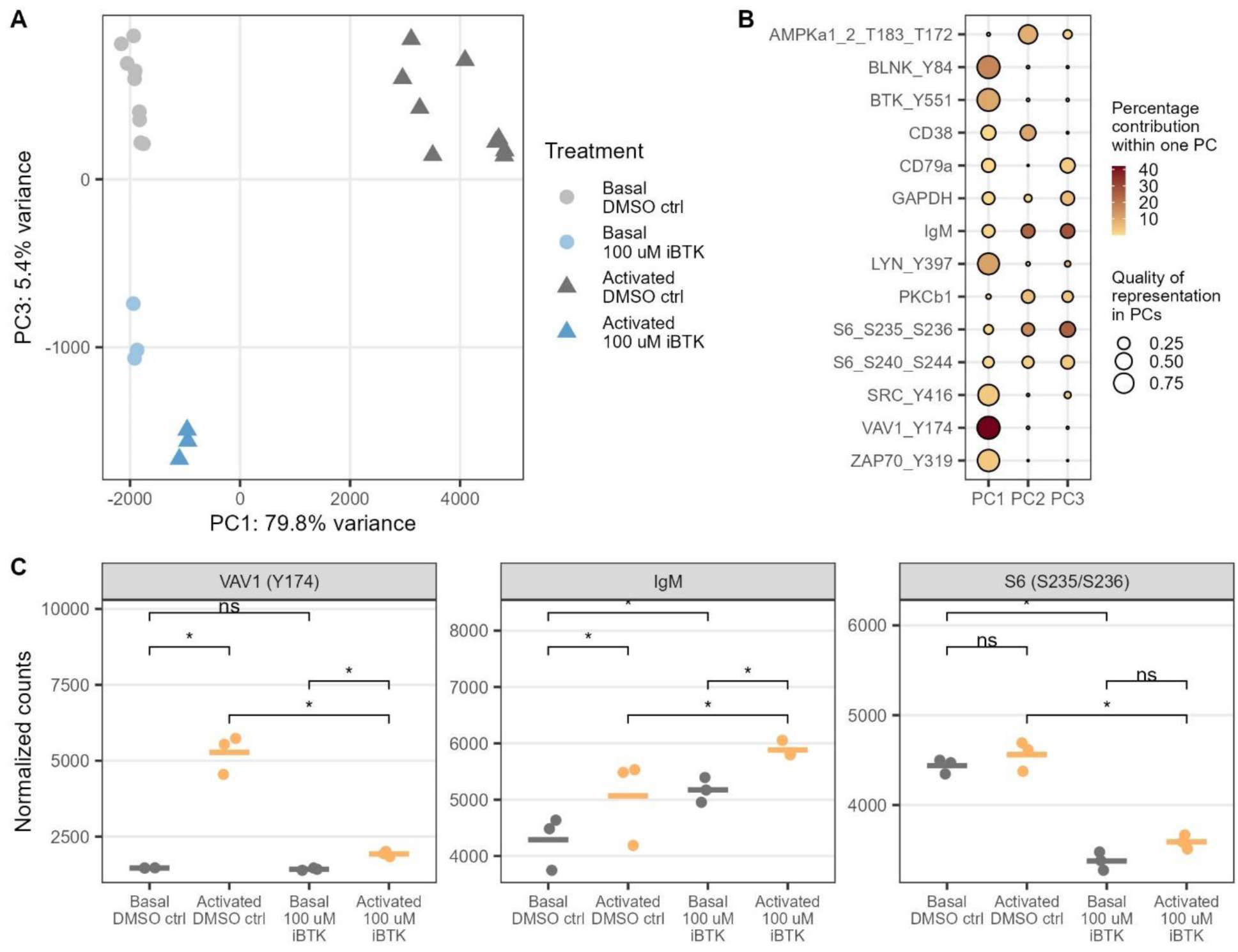
B-cell activation and inhibition by iBTK results in distinct signaling states. A. Principal component analysis (PCA) of basal, activated and 100 µM iBTK-inhibited cells. PCA was performed on all samples (normalized counts, Suppl. Fig. S6), selected samples were visualized. N = 3 biological replicates per condition. B. Proteins that contribute ≥ 2.5% to either PC1, PC2, or PC3. C. Proteins contributing most to PC1 (VAV1), PC2 (IgM), and PC3 (S6) as determined in Fig. 2B (individual replicates and mean normalized ID-seq count data). Statistical significance was determined with DEA (Fig 1C+D) using a Wald test with BH-adjusted p-value threshold ≤ 0.05.

PCA allowed us to extract which proteins contributed most to each PC and thus to the distinct signaling states. In agreement with the DEA, phospho-proteins contributing to PC1, capturing the differences between the basal and activated signaling states, included BLNK, BTK (Y551), LYN, and VAV1 (Fig. 2B). The proteins contributing to PC3, capturing the difference between the conditions with and without inhibitors, were CD79a, GAPDH, IgM, and phospho-S6. Analyzing the phosphorylation levels of the top contributing proteins to each PC sheds light on the PCA findings (Fig. 2C). VAV1 exhibited high phosphorylation (Y174) upon activation and reverted to basal levels upon treatment with iBTK (Fig. 2C left). Total IgM levels were slightly increased in the activated state and increased even more upon iBTK treatment (Fig. 2C middle). S6 (S235/S236) phosphorylation was strongly decreased by iBTK in both the basal and activated conditions, reaching new levels, and thereby illustrating how this phospho-protein especially contributed to the repressed signaling state (Fig. 2C right). To summarize, the core BCR phospho-proteins contributed to the activated signaling state, while more downstream phospho-proteins and non-phospho-proteins made the repressed signaling state different from the basal or activated states, indicating that different parts of this interconnected B-cell signaling network contribute to different overall outcomes or states of the network.

#### iBTK disrupts parts of the B-cell signaling network in a dose-dependent manner

To further understand how iBTK affects the change from basal to activated to repressed state, we included different doses of iBTK (10-fold increments from 0.1 µM to 100 µM). We then determined the effect of each iBTK dose on the signaling activation using DEA (Fig. 3A and Suppl. Fig. S8; log2(fold change) > log2(1.1), p-adjusted < 0.05). 15 phospho-proteins were significantly more abundant in the activated state compared to the basal state, and 14 of these 15 protein levels were significantly decreased by 100 µM iBTK treatment.

**Figure 3:**
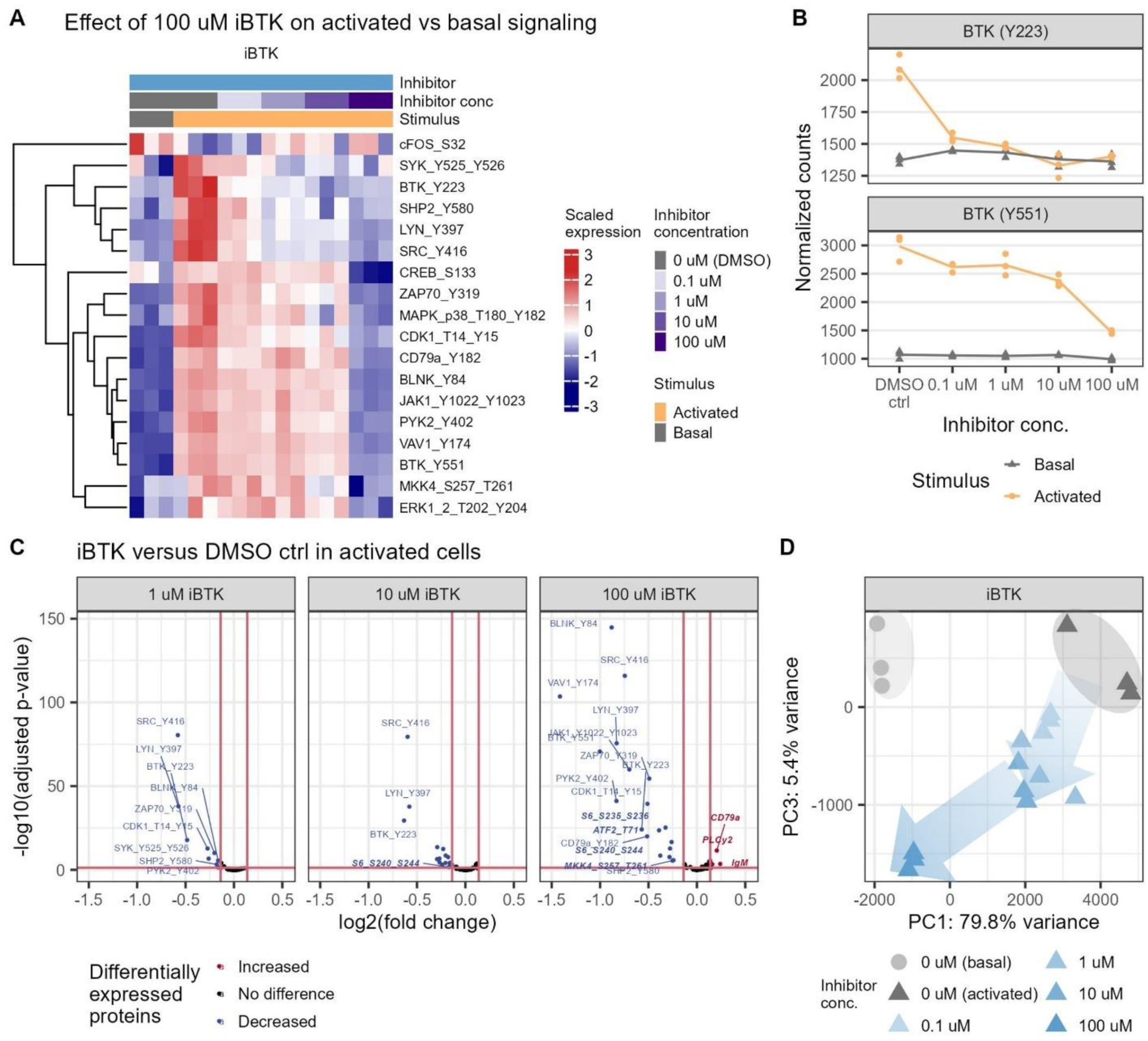
iBTK disrupts the entire B-cell signaling network and induces a dose-dependent repressed signaling state. N = 3 biological replicates per condition. A. Heatmap visualization of significant proteins from the DEA of activated versus basal treatment and the effect of iBTK dose on the activated versus basal comparison. Log2(fold change) threshold ≥ log2(1.1), BH-adjusted p-value threshold ≤ 0.05. B. BTK phosphorylation at Y223 and Y551 upon activating treatment and iBTK treatment (individual replicates and mean normalized ID-seq count data). Significance was determined in the DEA (see Fig. 3A and Suppl. Fig. S8). C. Volcano visualization of the DEA of iBTK dose compared to DMSO control in activated cells only (0.1 µM iBTK not shown). Log2(fold change) threshold ≥ log2(1.1), Wald test with BH-adjusted p-value threshold ≤ 0.05. Proteins uniquely identified in this analysis compared to A are highlighted in bold and italic. D. PCA of basal, activated and 0.1, 1, 10, and 100 µM iBTK-inhibited cells, with trend areas and direction indicated.

Hierarchical clustering of the significantly affected proteins revealed two main clusters that responded differently to iBTK: the top cluster of phospho-proteins was decreased by low doses of iBTK (1 µM and higher) and the bottom cluster was only significantly decreased by high doses (Fig. 3A). BTK phospho-site Y223 clustered together in the inhibitor sensitive cluster, together with LYN and SYK, while BTK phospho- site Y551 clustered in the less sensitive cluster, together with BLNK and VAV1. Indeed, the individual phospho-protein data showed a drastically different dose-dependent pattern of inhibition for the two BTK phospho-sites (Fig. 3B). Notably, BTK-Y223 is involved in the catalytic activity of BTK, and previous research has shown inhibition of BTK-Y223 by ibrutinib^22,46^.

Next, we focused on the activated signaling state only and compared (phospho-)protein levels in inhibitor-treated samples with the untreated DMSO control. This analysis identified any (phospho-)proteins that were significantly altered by iBTK treatment, regardless of their activation compared to basal signaling. For instance, phosphorylation levels of S6 (both phospho-sites) did not change upon activating treatment, and thus, no effect of iBTK on the activated versus basal signaling of S6 was observed (Fig. 3A). However, phosphorylation of S6 was significantly decreased upon iBTK treatment when comparing the iBTK treatment versus DMSO control in activated cells (Fig. 3C). Broadly, this analysis showed that higher iBTK doses led to increases in the number of proteins affected, the magnitude of the effect, and the significance level.

We then analyzed the PCA signaling state landscape to map along what axis iBTK propelled cells from the activated state without inhibitor towards the repressed state induced by high iBTK. Visual inspection of the intermediate iBTK conditions demonstrated that these samples map approximately equidistant between the activated and repressed state, and imply a repression trajectory that is slightly more along PC3 at low doses and then moves along PC1 at higher doses (Fig. 3D).

Together, we observed that the clinically approved and well-characterized inhibitor iBTK disrupts the entire B-cell signaling network, rather than only its target BTK and proteins downstream of BTK, even at low doses, and that this inhibition dose-dependently induces a repressed signaling state. These observations underscore that different pathways within the B-cell signaling network are tightly interconnected.

#### Network-wide signaling disruptions along different repression axes by iSYK and iNFκB

To corroborate whether other small molecule inhibitors also provoked network-wide signaling disruptions in DLBCL, we investigated iSYK, which is under investigation in clinical trials^33–35^, and iNFκB, which is a pre- clinical inhibitor^36,37^. In line with iBTK, both iSYK and iNFκB were able to disrupt activated B-cell signaling across the network (Suppl. Fig. S9). When the signaling network was inhibited maximally (100 µM), considerable overlap in signaling proteins affected by the different inhibitors was found (Fig. 4A), illustrating the high degree of crosstalk and connectivity in the network. However, at intermediate doses, we observed that iSYK (Fig. 4B) and iNFκB (Suppl. Fig. S10) presented subtle but definite differences in dose-dependent inhibition patterns compared to iBTK. For instance, DEA of the effect of iSYK doses on the activated versus basal signaling state, followed by heatmap visualization and hierarchical clustering revealed three main clusters (Fig. 4B): one strongly inhibited cluster, with SYK and PLCγ1, one weakly inhibited cluster even at high iSYK doses, which included BTK-Y551 and VAV1, and one moderately inhibited cluster (from 1 µM iSYK onwards), which included BTK-Y223 and LYN. Noteworthy, JAK1 clustered together with BTK-Y223 under iSYK treatment, but clustered together with BTK-Y551 under iBTK treatment. Indeed, the protein data confirmed that JAK1 phosphorylation was decreased only by the highest dose of iBTK, steadily decreased by increasing doses of iSYK, and decreased equally by 0.1, 1 and 10 µM iNFκB (Fig. 4C). Likewise, phospho-proteins CD79a, IKKa/b and MAPK p38 highlight that these three inhibitors demonstrated markedly different dose-dependent effects on phospho-protein levels at intermediate doses (Suppl. Fig. S11).

**Figure 4:**
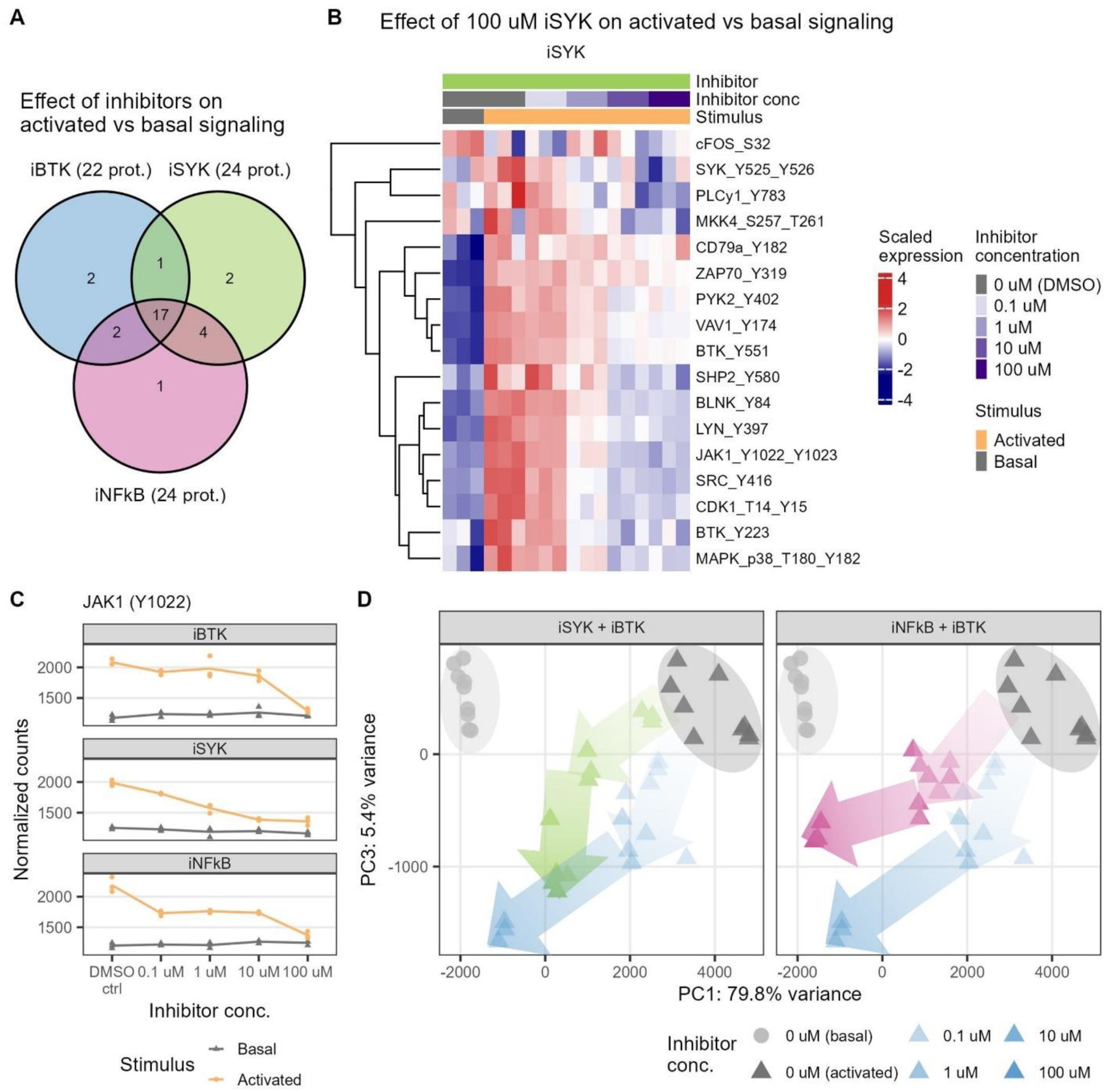
Corroboration of network-wide signaling disruptions with iSYK and iNFκB. N = 3 biological replicates per condition. A. Number of proteins that are significantly affected in their activation (activated versus basal) by any dose of an inhibitor. Significance was determined via DEA with a log2(fold change) threshold ≥ log2(1.1) and a Wald test with BH-adjusted p-value threshold ≤ 0.05. B. Heatmap visualization of significant proteins from the DEA of activated versus basal treatment and the effect of iSYK dose on the activated versus basal comparison. Log2(fold change) threshold ≥ log2(1.1), Walt test with BH-adjusted p-value threshold ≤ 0.05. C. JAK1 phosphorylation (Y1022) upon activation and inhibition (individual replicates and mean normalized ID-seq count data). Significance was determined in DEA (see Fig. 3A, Fig. 4B, and Suppl. Fig. S10). D. PCA of basal, activated and iSYK- or iNFκB-inhibited cells. iBTK samples (same as in Fig. 3D) were visualized for comparison. Trend areas and directions are indicated with arrows.

Since all three inhibitors demonstrated the ability to disrupt the entire B-cell network at high doses but exhibit varying effects at intermediate doses, we explored how iSYK and iNFκB treatment acted within the PCA signaling state landscape compared to iBTK. We observed that iSYK, and to a lesser extent iNFκB, at high doses indeed pushed the cells towards the repressed state (Fig. 4D). Furthermore, treatment with low and intermediate doses of iSYK pushed cells along a PC1-PC3-oriented axis towards the repressed state, while iBTK treatment pushed cells along a PC3-PC1-oriented axis. iNFκB treatment pushed cells mostly along a PC1-oriented axis, and fully iNFκB-inhibited samples resembled the basal state more than iBTK or iSYK inhibited samples. Taken together, the inhibitors repressed the entire B-cell signaling network along distinct repression axes within the signaling state landscape.

#### Combinatory inhibitor treatments effectively induce repressed signaling states

Inspired by the work of Kholodenko and co-workers who demonstrated how cells states could be transformed along state transition vectors^15,16,47^, we hypothesized that combinatory treatment at low doses would also propel cells towards the repressed signaling state. Indeed, some synergistic effects have been described for iBTK and a PI3K inhibitor^48,49^, and for iSYK and a LYN inhibitor^50^. Thus, combinatory treatment could result in more effective disruptions of the B-cell signaling network than high doses of the single inhibitors. To test this hypothesis, we analyzed the (phospho-)protein responses with ID-seq after treating DLBCL cells with different combinations of iBTK, iSYK, and iNFκB.

First, we mapped the entire signaling state landscape of this experiment with PCA (Suppl. Fig. S12). In line with the single inhibitor experiment, PC1 captured the main differences between the basal and activated signaling states and PC2 captured the differences between the basal state and the repressed state that is induced by combining all three inhibitors. Moreover, BLNK, BTK (Y551), and VAV1 contributed most to the difference between the basal and activated states, while BLNK, IgM, and LYN contributed most to reaching the three-inhibitor repressed state (Suppl. Fig. S13).

As reference, we treated cells with 10 µM of the single inhibitors. In the signaling state landscape, these conditions mapped approximately equidistant between the basal and activated states (Fig. 5A, Suppl. Fig. S14). Subsequently, we studied the effect of combining two inhibitors, each at 2.5 µM. The samples treated with iBTK + iSYK and with iBTK + iNFκB clustered closely together with the 10 µM iSYK treated samples (Fig. 5B). The iSYK + iNFκB combination was slightly shifted compared to the single inhibitor treated samples, suggesting stronger inhibition by this combinatory treatment. When we combined the three inhibitors together at equal doses, we observed an even larger shift in the signaling state landscape, in the extended direction as seen with the two-inhibitor treated cells (Fig. 5C). As expected, higher doses of the inhibitors induced a more repressed state.

**Figure 5:**
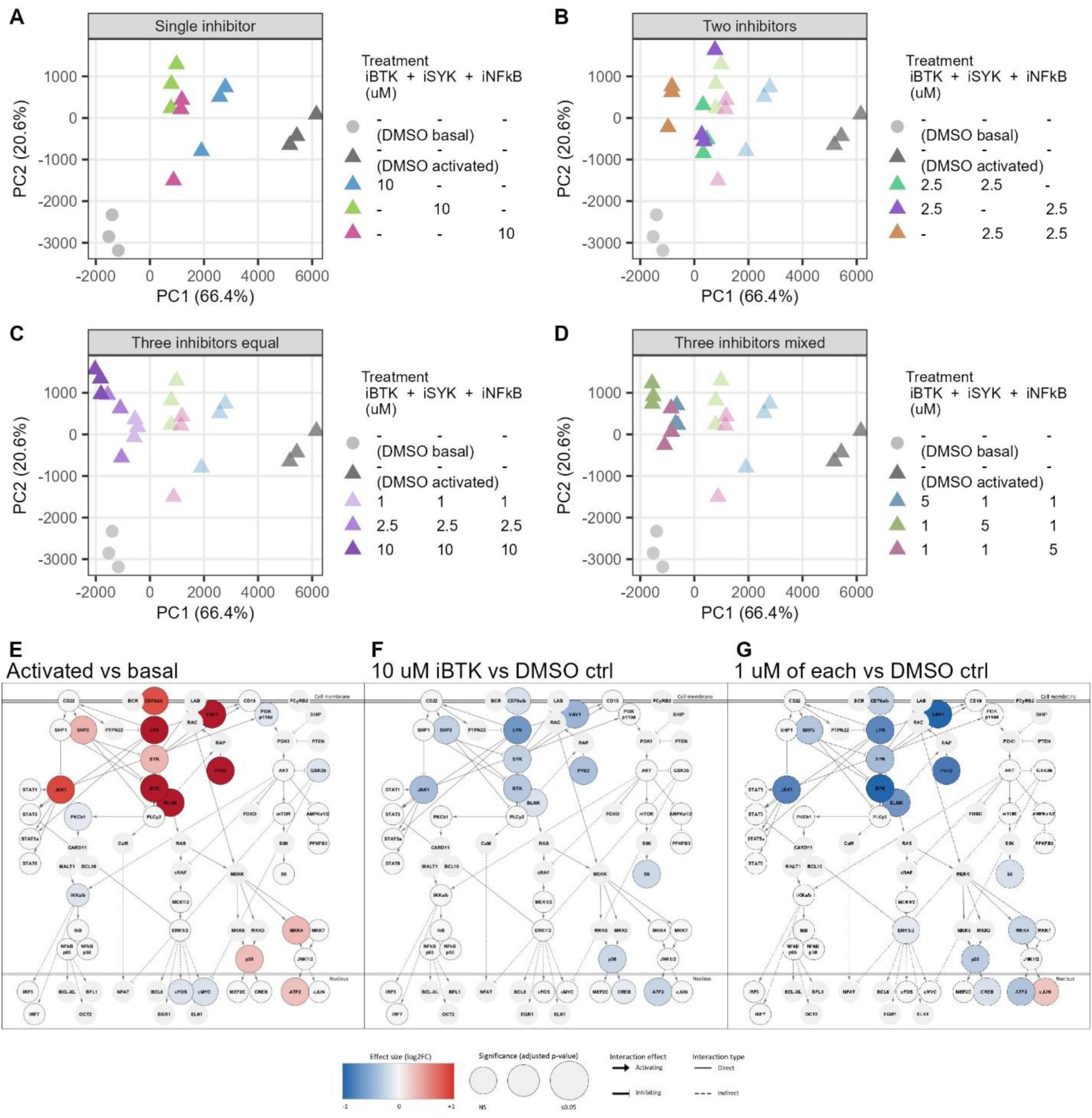
Inhibitor combination treatments effectively induce repressed signaling states. N = 3 biological replicates per condition. A-D. PCA of basal, activated, and (combinatory) inhibited cells. PCA was performed on all samples (Suppl. Fig. S12), and selected conditions were visualized. DMSO controls (grey) and single inhibitor conditions (blue, green and pink) are visualized in all panels for comparison. E-G: Network visualization of DEA of activated cells compared to basal cells (E), and of inhibitor treatment compared to DMSO control (10 µM iBTK in F, 1 µM of iBTK + iSYK + iNFκB in G). Walt test with BH-adjusted p-value threshold ≤ 0.05. See Suppl. Fig. 1 for a larger schematic of the network in E-G.

The above-described effects of combining two or three inhibitors at equal doses on the signaling state prompted the question if we can manipulate how the cells move through the signaling state landscape by combining the three inhibitors at mixed doses. We tested one inhibitor at 5 µM while keeping the other two inhibitors at 1 µM (Fig. 5D). As expected, the mixed three-inhibitor conditions mapped in between the conditions with 1 µM or 10 µM of each inhibitor. Moreover, we observed the strongest repression when iSYK was added at 5 µM, compared to when iBTK or iNFκB were added at 5 µM, which is in line with the two-inhibitor treatment (Fig. 5B). Thus, we found that low dose combinations of iSYK, iBTK and iNFκB together were more effective in inducing a repressed signaling state than 10 µM of a single inhibitor.

To place the combinatory inhibitor results in the context of the B-cell signaling network, we performed DEA, comparing the basal and activated states, and the effect of the inhibitor treatments on the activated state. As shown before, the activated state consisted of network-wide phosphorylation events, especially in the core BCR signaling hub (Fig. 5E). Treatment of the activated state with 10 µM iBTK alone resulted in substantial inhibition of the network, mainly in the core BCR hub, but also in MAPK p38 signaling and JAK1 signaling (Fig. 5F). Strikingly, combinatory treatment with 1 µM of iBTK + iSYK + iNFκB disrupted the signaling network more strongly than the single inhibitor treatments (Fig. 5G, Suppl. Fig S15). Individual protein data exemplified either strong repression across all inhibitor combinations that were investigated, or inhibitor-dependent differences in the strength of repression (Suppl. Fig. S16). Together, combinatory treatment with low doses of multiple inhibitors was more effective in disrupting activated B-cell signaling in these malignant B-cells.

## Discussion

Malignant cells can hijack signaling pathways for their growth, which complicates defining the effects of drugs. We have shown that ID-seq can be a powerful tool to capture the phosphorylation state of signaling networks, and study cellular signaling in the context of larger networks. Using a panel of 111 antibodies, we uncovered major cellular signaling state changes in aggressive B-cell lymphoma (ABC-DLBCL) in the context of activation (Fig. 6A+B) and BTK inhibition (Fig. 6C). Decreased phosphorylation of the core BCR signaling hub was observed, where also proteins that are classically depicted upstream of BTK were inhibited. Moreover, inhibition of the activated signaling state did not revert to the basal state, but instead induced a new, repressed state. Exploring two other therapeutic inhibitors, iSYK and iNFκB, confirmed that these inhibitors perform equally well regarding the disruption of the entire B-cell signaling network and propelling cells towards a repressed signaling state, although each inhibitor had a unique axis of repression. When we applied combinatory inhibitor treatments, thus pushing the network from multiple angles, we observed strong inhibitory effects with low doses (Fig. 6D). These findings show that cellular signaling occurs in complex networks, and underscore how quantification of the signaling state landscape provides insights into how combination drug treatments can bring cells in a desired network state.

**Figure 6:**
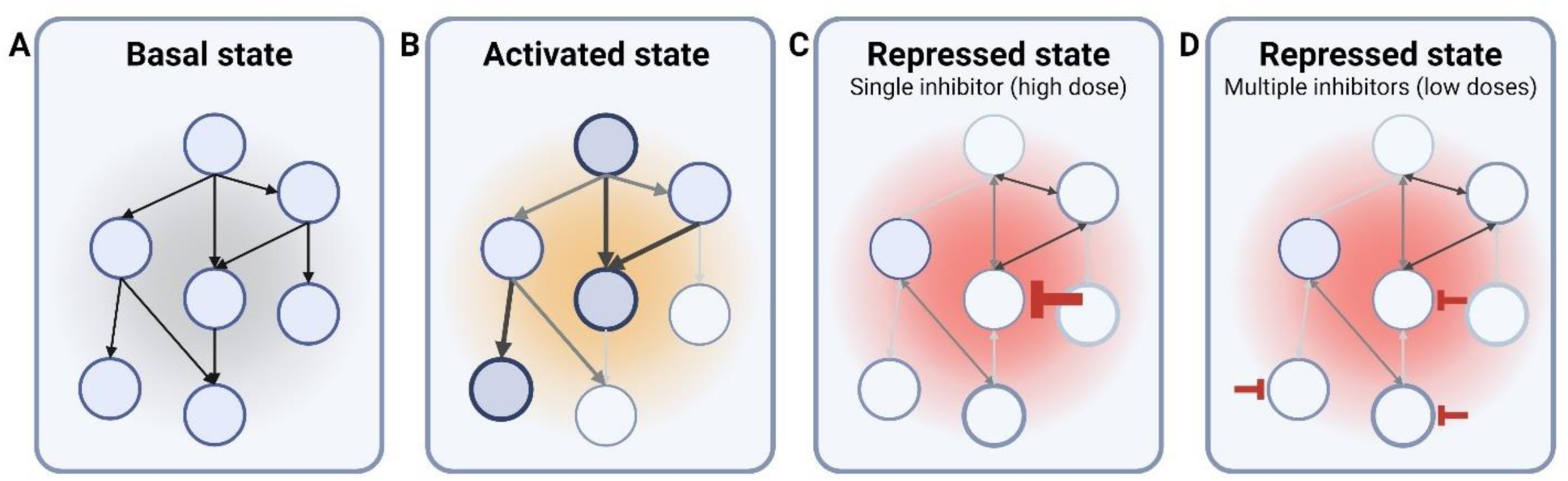
Inhibition of the B-cell network results in a transition from an activated signaling state to a repressed state. A. Basal state of a signaling network. B. Activated state. C. Repressed state induced by a single inhibitor at a high dose. D. Repressed state of a signaling network induced by combinatory inhibitor treatment at low doses.

A major advantage of ID-seq is the vast number of targets that can be measured. In this study, we have used a panel of 111 antibodies, broadly covering the B-cell network and additional pathways of interest, and this panel could be expanded even further. In addition, the ID-seq method can be expanded towards single-cell measurements^51^, combined with RNA-seq measurements^52,53^, and has the potential to be integrated with commercial kits due to the antibody barcode design. ID-seq and other large-scale high-throughput approaches are key in transitioning from a linear cascade view of cellular signaling to an integrated network view. For instance, our work elucidated that iBTK treatment resulted in two sets of proteins that had different dose-response curves to the inhibitor, suggesting two interconnected modules in the signaling network that are regulated somewhat separately. The two measured phospho-sites of BTK (Y223 and Y551) clustered in the different sets, hinting at a regulatory mechanism that works through BTK phosphorylation. BTK-Y551 is phosphorylated by LYN and plays a role in BTK activation, while BTK-Y223 is an autophosphorylation step and essential for the catalytic activity^46^, and iBTK (ibrutinib) has been described to reduce Y223 phosphorylation^22^. It is also known that BTK is a key signaling protein in B-cell signaling, connecting different pathways, such as chemokine, integrin, TLR and Fc receptor signaling, to BCR signaling^11,54^. These results confirm an important role for BTK, which should be explored further.

The observation that iBTK, iSYK and iNFκB all caused major disruptions of activated B-cell signaling and induced a repressed signaling state was striking. As SYK is located in proximity to BTK in the B-cell signaling network, both physically and functionally^41,54^, we expected that inhibiting these two proteins would have comparable effects. In addition, both proteins are located relatively upstream in the B-cell signaling network. NFκB, on the other hand, is located much more distally in the network^55,56^, and yet, our data revealed many changes in proximal B-cell signaling upon NFκB inhibition. iNFκB is a preclinical drug, described to block NFκB transcriptional activity^36,37^, although the exact mechanism is not fully elucidated. Although off-target effects of iNFκB are a possibility, this has not been described in previous studies on iNFκB^57–59^, and we observed significant effects at low doses. For these reasons, we believe that these findings illustrate the interconnectivity of the B-cell signaling network. Indeed, many reverse connections and feedback loops have been reported in the B-cell signaling network: reverse interactions between LYN and SYK^60–62^, an autoactivation loop on SYK^13^, interactions between PDK1 and MALT (PI3K branch and NFκB branch)^10^, and various inhibitory phosphatase interactions^63^, and there could be many more. The question then arises whether it is necessary that an inhibitor is specific for one target, or whether it is more important that the overall desired effect on the signaling network is achieved. As also proposed by other researchers, drug development research might benefit from focusing on the signaling state effects of inhibitors rather than specific protein inhibition^14,15^.

The signaling state landscape of iBTK, iSYK and iNFκB described distinct axes of repression for these inhibitors across a concentration range. These different axes of repression, together with the different dose-response patterns between sets of proteins, suggest a balancing act of B-cell signaling proteins that can be shifted towards a more activated or repressed state from different angles by different inhibitors. The development of new and improved treatment options for DLBCL is a research topic still requiring attention since many patients do not respond to or relapse with chemo(immuno)therapy^17,18^. Previous work on combinatory treatment for DLBCL has either focused on the clinical effects of combining a small molecule inhibitor with standard chemo(immuno)therapy^28,30^ or on combining ibrutinib (iBTK) with other small molecule inhibitors in viability screens^48,64^. A few studies have investigated other small molecule inhibitor combinations, such as BTK + PI3K inhibition^49^, LYN + SYK inhibition^50^, or PI3Kδ + PI3Kα inhibition^65^, with promising results and synergistic effects at the molecular level. Our findings on two- and three- inhibitor combinatory treatments offer additional insights in drug effects at the signaling level.

In conclusion, we have demonstrated how quantification of the B-cell signaling state allows us to explore the interconnectivity of signaling networks. Our findings indicate that B-cell signaling acts as a network rather than a linear cascade and highlight the potential for combination treatment to obtain effective results using low doses, thus opening up avenues for more effective therapies with fewer side effects.

## Methods

### Materials

All reagents and materials are listed in Table 1.

**Table 1:** Key resources table. NA = not applicable.

| Reagent or resource | Source | Identifier |
| --- | --- | --- |
| Antibodies |  |  |
| Antibodies used for ID-seq are listed in Suppl. Table S1 | See Suppl. Table S1 | NA |
| Antibodies used for flow cytometry are listed in Suppl. Table S2 | See Suppl. Table S2 | NA |
| Chemicals, peptides, and recombinant proteins |  |  |
| 4x Laemmli Sample Buffer | BioRad | Cat#1610747 |
| 10x Tris/Glycine/SDS electrophoresis buffer | BioRad | Cat#1610772 |
| alg: AffiniPure F(ab')<sub>2</sub> Fragment Goat Anti-Human IgA + IgG + IgM (H+L) | Jackson ImmunoResearch | Cat#109-006-064 |
| Bovine serum albumin (BSA) | Sigma | Cat#A4503-50G |
| Dibenzocyclooctyne-S-S-N-hydroxysuccinimidyl ester (DBCO-S-S-NHS) | Sigma | Cat#761532 |
| Dimethyl sulfoxide (DMSO) | Sigma | Cat#276855-100mL |
| 10 mM Deoxynucleotide (dNTP) Solution Mix | New England Biolabs | Cat#N0447L |
| DL-Dithiothreitol (DTT) solution | Sigma | Cat#43816 |
| 1x Dulbecco's phosphate buffered saline (DPBS) | Gibco | Cat#14190-094 |
| 0.5M EDTA solution, pH 8.0 | Lonza | Cat#51201 |
| Fetal bovine serum (FBS) | Gibco | Cat#A3160801 |
| FcR Blocking Reagent Human | Miltenyi Biotec | Cat#130-059-901 |
| Hydrogen peroxide (H<sub>2</sub>O<sub>2</sub>) solution, 35% | Thermo Fisher Scientific | Cat#202460010 |
| iBTK: Ibrutinib (PCI-32765, Imbruvica) | Selleckchem | Cat#S2680 |
| iNFκB: QNZ (EVP4593) | Selleckchem | Cat#S4902 |
| iSYK: R406 | Selleckchem | Cat#S2194 |
| Klenow Fragment (3'→5' exo-) | New England Biolabs | Cat#M0212L |
| 1M Magnesium chloride (MgCl<sub>2</sub>) | Thermo Fisher Scientific | Cat#AM9530G |
| Nuclease-free (NF) water | Ambion | Cat#AM9937 |
| Paraformaldehyde (PFA) | Merck | Cat#1040051000 |
| Penicillin/streptomycin | Gibco | Cat#15140-122 |
| Phosphate buffered saline (PBS) | Gibco | Cat#14190-094 |
| Precision Plus Protein™ All Blue Prestained Protein Standards | BioRad | Cat#1610373 |
| Q5® High-Fidelity DNA Polymerase | New England Biolabs | Cat#M0491S |
| Q5® Reaction Buffer | New England Biolabs | Cat#B9027S |
| RPMI Medium 1640 | Gibco | Cat#52400-025 |
| Sodium chloride (NaCl) | Sigma | Cat#S3014-500G |
| Sodium hydrogen carbonate (NaHCO<sub>3</sub>) | Merck | Cat#1.06329.1000 |
| Sodium azide (NaN<sub>3</sub>) | Sigma | Cat#S2002 |
| UltraPure™ Salmon Sperm DNA Solution | Invitrogen | Cat#15632011 |
| 1M Tris-HCl, pH 7.5 | Gibco | Cat#15567-027 |
| Triton X-100 Surfact-Amps detergent solution | Thermo Fisher Scientific | Cat#85112 |
| Critical commercial assays |  |  |
| Agilent High Sensitivity DNA kit | Agilent Technologies | Cat#5067-4626 |
| KAPA HiFi HotStart Ready mix | Rosh | Cat#KK2601 |
| Qubit dsDNA High Sensitivity Assay kit | Thermo Fisher Scientific | Cat#Q32851 |
| Zymo DNA Clean & Concentrator-5 purification kit | Zymo Research | Cat#D4014 |
| Deposited data |  |  |
| Raw and analyzed ID-seq data | This paper | [GSE279584](https://www.ncbi.nlm.nih.gov/geo/query/acc.cgi?acc=GSE279584) |
| Experimental models: Cell lines |  |  |
| Human: HBL1 | Supplied by Dr. B. Scheijen (Dept. Pathology, Radboudumc) | RRID:CVCL_4213 |
| Oligonucleotides |  |  |
| DNA oligos used for antibody barcodes are listed in Suppl. Tables S1 and S3 | Biolegio, The Netherlands | NA |
| Primers and other oligos used for library preparation are listed in Suppl. Tables S3, S4, and S5 | Biolegio, The Netherlands | NA |
| Software and algorithms |  |  |
| BD FACSuite software v1.0.6 | BD Biosciences | NA |
| R version 4.3.3 | The R Foundation | https://www.r-project.org/ |
| RStudio | RStudio, USA | https://www.rstudio.com/ |
| BioRender | BioRender, Canada | https://biorender.com/ |
| Other |  |  |
| 4–15% Mini-PROTEAN™ TGX Stain-Free™ Protein Gels | BioRad | Cat#4568086 |
| 96-well Clear V-Bottom Not Treated Polypropylene Storage Microplate | Falcon | Cat#353263 |
| HTS multiwell storage plate, 96-well, u-bottom, without lid, polypropylene | Falcon | Cat#351196 |
| Multiplate PCR plates 96-well clear | BioRad | Cat#MLP9601 |
| Ultracel-100 regenerated cellulose membrane, 0.5 mL sample volume | Millipore | Cat#UFC510096 |
| Zeba Spin Desalting Columns, 0.5ml 40K MWCO | Thermo Fisher Scientific | Cat#87767 |

#### Cell culture

HBL1 cell line (RRID: CVCL_4213) was used as model for B-cell lymphoma (ABC-DLBCL). This cell line is EBV-negative, was established from a Japanese 65-year-old male in 1988^66^, and has been STR validated. Cells were cultured in RPMI 1640 medium supplemented with 10% (v/v) fetal bovine serum (FBS) and 1% (v/v) penicillin/streptomycin (P/S) at a density of 0.20-2.0*10^6^ cells/mL in a humidified incubator at 37°C with 5% CO_2_.

#### B-cell preparation and treatment

Before experiments, cells underwent a medium change to serum-poor medium (RPMI 1640 medium with 2% FBS and 1% P/S) and were allowed to rest for 60 min at 37°C (at 3.3*10^6^ cells/mL). Inhibitors of SYK (R406, referred to as iSYK), BTK (ibrutinib, referred to as iBTK), and NFκB (QNZ, referred to as iNFκB) were dissolved in DMSO. 0.5*10^6^ cells per well were distributed in 96-well plates, inhibitors were added at 0, 0.1, 1, 10, 100 µM final concentrations (keeping the DMSO content at 1%), and incubated for 60 min at 37°C with 5% CO_2_. For combination treatments, the respective combinations of inhibitors were prepared in separate tubes first and then distributed in 96-well plates, keeping the final DMSO content at 1%. Both 0 µM with DMSO and 0 µM with PBS were taken along as controls. Then, the stimulus was added, either PBS or 15 µg/mL F(ab’)₂ Fragment Goat Anti-Human IgA + IgG + IgM (H+L) (anti-Ig) + 15 mM hydrogen peroxide (H_2_O_2_) final concentrations, and incubated for 20 min at 37°C with 5% CO_2_. Sample fixation with 4% paraformaldehyde (PFA) was performed for 15 min at room temperature (RT) and was stopped via centrifugation for 5 min at 1500 rcf and resupination in phosphate buffered saline (PBS). Samples were split into two aliquots (0.25*10^6^ cells each) and transferred to 96-well plates to facilitate measurements with both phospho flow cytometry and immunodetection by sequencing (ID-seq).

#### Phospho flow cytometry of fixed cells

The phospho flow cytometry measurements and data analysis were performed as described previously by Witmond *et al.* (2024)^45^. Briefly, PFA-fixed cells were centrifuged, permeabilised with 0.1% Triton X- 100 in water, washed 1x with FACS buffer (Dulbecco’s PBS with 0.1% bovine serum albumin [BSA], 0.05% NaN_3_ and 0.5 mM EDTA), and stained with a panel of fluorescently labelled antibodies (Suppl. Table S2). Cells were washed 3x and resuspended in FACS buffer for measurements on the BD FACSVerse (BD Biosciences) (50000 events per sample). Compensation matrices created from single-stain controls of each antibody were applied to all measurements. We refer to Witmond *et al.* (2024) for more details on the methodology.

Flow cytometry data were analysed in RStudio (see session information in the analysis files for all packages). Gating was performed using the flowCore^67^ and ggcyto^68^ packages. Debris was removed first (FSC-A vs SSC-A), then singlet cells were selected (FSC-H vs FSC-W), and then live cells were selected (FSC-A vs active caspase 3 + cleaved PARP, BV421-A) (Suppl. Fig. S17). The gated data were processed further, calculating the median fluorescence intensity (MFI) per target, and visualized using the tidyverse package^69^. Significant differences between conditions were determined with the Kruskal-Wallis test (p < 0.001) and post-hoc Dunn’s test with Benjamini-Hochberg (BH)-correction for multiple testing (p-value threshold ≤ 0.05).

#### Conjugation of DNA oligos to antibodies

The functionalisation and conjugation of 50 µg antibody to 5’ azide modified DNA oligos (Ab barcodes) via NHS click chemistry was based on the work of Van Buggenum *et al.* (2016 and 2018)^20,70^ and Rivello *et al.* (2021)^53^. All antibodies used for ID-seq were purchased carrier-free (Suppl. Table S1). See Suppl. Fig. S18 for the design of the Ab barcodes, Suppl. Table S1 for target-specific barcode sequences, and Suppl. Table S3 for the complete DNA oligo sequence.

Briefly, antibodies were first buffer exchanged to 0.2 M sodium bicarbonate (NaHCO_3_) buffer (pH 8-9) with Zeba™ Spin Desalting Columns. 15x molar excess dibenzocyclooctyne-S-S-N-hydroxysuccinimidyl ester (DBCO-S-S-NHS) in DMSO was added and incubated for 2 hours at RT in the dark. Antibodies were again buffer exchanged to 0.2 M NaHCO_3_ buffer with Zeba™ Spin Desalting Columns to remove excess DBCO-S-S-NHS. Then, Ab barcode was added in 3x molar excess, incubated for 1 hour at RT in the dark, followed by overnight incubation at 4°C. Storage buffer, containing 0.05% NaN_3_ and 1 mM EDTA in DPBS, was added and antibodies were aliquoted and stored at -20°C. Labelling efficiency was determined by non-reducing SDS-PAGE gel electrophoresis; 0.5 µg antibody in 1x Laemmli sample buffer was loaded on a 4–15% Mini- PROTEAN™ TGX Stain-Free™ protein gel. Gels were imaged on a GelDoc XR+ imaging system (BioRad; see Suppl. Fig. S19 for representative conjugation results). To create the ID-seq antibody panel, aliquots of each DNA-labelled antibody were defrosted, antibodies were combined in equal amounts, and the panel was concentrated in storage buffer with the Amicon 100K Ultra Centrifugal Filter Unit according to the manufacturer’s protocol.

#### Immunodetection by sequencing (ID-seq) of fixed cells

The immunodetection by sequencing (ID-seq) of fixed cells was based on the work of Van Buggenum *et al.* (2018)^20^ with some modifications to optimise the procedure for B-cell lines. Briefly, PFA-fixed cells in 96-well plates (0.25*10^6^ cells/well) were centrifuged for 5 min at 1500 rcf and permeabilised for 10 min in NF water with 100 mM Tris-HCl, 150 mM NaCl and 0.1% Triton X-100 at RT. Samples were then blocked for 30 min at RT with blocking buffer (DPBS with 3% BSA, 20 µg/mL single-stranded salmon sperm DNA, 0.1% Triton X-100, 1/50 FcR blocking reagent). After centrifugation, cells were resuspended in 30 µL staining solution, consisting of 0.15 µg/mL of each DNA-labelled antibody in blocking buffer (see Suppl. Table S1 for all antibody information), and incubated overnight at 4°C. Cells were washed 6x with 3% BSA in DPBS, consisting of 4 wash steps with immediate centrifugation and 2 wash steps with 15 min incubation at RT before centrifugation. At the last wash step, samples were transferred to a new 96-well plate, resuspended in 50 µL release buffer, consisting of 30 mM dithiothreitol (DTT) in 200 mM NaHCO_3_ (pH 8-9) and incubated for at least 1.5 hours to release the antibody barcodes (Ab barcodes) from the antibodies. Cells were then centrifuged for 5 min at 1500 rcf and the supernatant with the released Ab barcodes was collected.

To prepare ID-seq libraries, well-specific barcodes for each plate were attached to the released Ab barcodes by mixing 9 µL released Ab barcodes with 0.5 µM final concentration well barcode (all complete barcode and primer sequences are listed in Suppl. Table S3, see Suppl. Table S4 for well-specific barcodes) and incubated for 30 min at 25°C. To elongate the annealed well barcodes and Ab barcodes, a polymerase reaction was performed with Klenow Fragment (3′→ 5′ exo-). 1.5 mM dNTP, 0.15 units/µL Klenow Fragment and 7.5 mM MgCl_2_ (final concentrations) was added per well and incubated for 10 min at 21°C, followed by 60 min at 37°C and a final 10 min at 21°C in a T100 Thermal Cycler (BioRad). All samples were pooled, purified with the Zymo DNA Clean & Concentrator-5 purification kit according to the manufacturer’s protocol and eluted in NF water. The concentration of ID-seq libraries was measured after each purification step with the Qubit dsDNA High Sensitivity Assay kit according to the manufacturer’s protocol.

Then, an amplification PCR was performed. One reaction contained approximately 5 ng input DNA, 1x Q5 Reaction Buffer, 200 µM dNTP, 0.5 µM forward primer, 0.5 µM reverse primer, 0.02 units/µL Q5 High-Fidelity DNA polymerase, and NF water (50 µL total volume). The PCR program on a T100 Thermal Cycler (BioRad) consisted of 3 min at 98°C followed by 10 cycles of 15 sec at 98°C, 20 sec at 68°C, and 60 sec at 72°C, and finished with 60 sec at 72°C. The PCR product was purified with the Zymo purification kit and eluted in NF water.

An index PCR was performed to attach Illumina sequencing adapters. One reaction contained approximately 4 ng input DNA, 1x KAPA HiFi HotStart Ready mix, 0.3 µM Nextera forward primer (Suppl. Table S3), 0.3 µM unique indexing reverse primer (containing a Nextflex 8bp index, see Suppl. Table S5), and NF water (25 µL total volume). The PCR program on a T100 Thermal Cycler (BioRad) consisted of 45 sec at 98°C followed by 5 cycles of 20 sec at 98°C, 30 sec at 64°C, and 20 sec at 72°C, and finished with 60 sec at 72°C. Finally, the PCR product was purified with the Zymo purification kit and eluted in NF water. ID-seq library sizes and quality were checked on a 2100 BioAnalyzer (Agilent) with the Agilent High Sensitivity DNA kit according to the manufacturer’s protocol. Paired-end sequencing of the ID-seq libraries was performed on an NextSeq500 sequencer (Illumina) with a target sequencing depth of approximately 1500 reads per antibody per sample.

#### ID-seq data analysis

The ID-seq sequencing data were demultiplexed with bcl2fastq software and the quality was assessed with MultiQC (version 1.9)^71^. Then, all reads were pseudo-aligned to the correct barcodes (Ab barcode and well barcode) and count tables were generated using kallisto^72^ and BUStools^73^. All count tables were analysed in RStudio with various packages (see session information in the analysis files for all packages). The quality of the data was assessed based on count distribution across wells, treatment conditions and target proteins. Data were normalised to account for cell number differences between samples with a geometric mean scaling factor: first, the geometric mean was calculated for each target and then the median of the target scaling factors was calculated for each sample. As the obtained count tables are similar to bulk RNA-seq data, differential expression analysis (DEA) comparing conditions of interest was performed on the unnormalized counts with the DESeq2 package^74^. The DEA returned log2 fold changes (log2(fold change)) of compared conditions as well as Wald test p-values and Benjamini-Hochberg (BH) adjusted p-values, where adjusted p-values <0.05 were considered statistically significant. Visualizations of differential expression results projected on the BCR signaling network were created with Cytoscape version 3.10.1^75^. Principal component analysis (PCA) with 5 principal components was performed on all samples within one experiment using the FactoMineR package^76^. Significant differences between specific conditions were determined with the Kruskal-Wallis test (p < 0.001) and post-hoc Dunn’s test with BH-correction for multiple testing (p-value threshold ≤ 0.05). Additional data processing and visualization was mainly done through the tidyverse package^69^ and a readable html documentation was generated with workflowR^77^.

## Data and code availability

All data analysis was performed using custom R scripts. These are publicly available as of the date of publication on GitHub: <u>huckgroup/DLBCL_manuscript</u>. The raw and processed ID-seq data has been deposited in NCBI’s Gene Expression Omnibus (GEO) under the accension number <u>GSE279584</u>. Any additional information required to analyse the data reported in this paper is available from the corresponding author upon request.

## Acknowledgments

This research was funded in part by Radboud University, the Dutch Research Council Spinoza Grant (WTSH), and the Dutch Research Council VENI Grant VI.Veni.202.228 (JAGLvB). AvS is supported by the Netherlands Organization for Scientific Research (NWO): the Institute of Chemical Immunology (project ICI00023), ZonMW (project 09120012010023), the Dutch Cancer Society (projects 12949 and 14726), and the European Research Council: Consolidator Grant (project 724281) and Proof-of-Concept Grant (project 101112687). Schematic illustrations were created in BioRender: Huck, W. (2024).

## Author contributions

Conceptualization: MW, ABvS, JAGLvB, WTSH

Methodology: MW, BB

Investigation: MW, PH

Visualization: MW

Funding acquisition: JAGLvB, WTSH

Project administration: MW

Supervision: JAGLvB, WTSH

Writing – original draft: MW, JAGLvB

Writing – review & editing: MW, ABvS, JAGLvB, WTSH

## Competing interests

Authors declare that they have no competing interests.

## Supplementary materials

Figure S1: Detailed schematic of the B-cell signaling network, with phospho-proteins measured in ID-seq panel indicated.

Figure S2: Increased phosphorylation upon treatment with both anti-Ig and H_2_O_2_ combined compared to treatment with anti-Ig or H_2_O_2_ alone.

Figure S3: Schematic overview of the experimental workflow of ID-seq.

Figure S4: Phospho flow cytometry measurements of CD79a (Y182) shows high concordance with ID-seq results.

Figure S5: No significant changes in (phospho-)protein levels upon treatment with DMSO compared to PBS.

Figure S6: Complete PCA signaling state space of the inhibitor concentration experiment.

Figure S7: Principal components PC1-3 visualized for DMSO controls (basal and activated) and 100 µM iBTK treatment.

Figure S8: Effect of iBTK dose on the comparison between activated and basal signaling.

Figure S9: Inhibition effect of 100 µM iSYK or 100 µM iNFκB versus DMSO control in activated cells.

Figure: S10: Effect of iNFκB concentration on the comparison between activated and basal signaling.

Figure S11: Different dose-response patterns in phosphorylation levels of CD79a, IKKa/b, and MAPK p38 upon activation and inhibition by iBTK, iSYK, and iNFκB.

Figure S12: Complete PCA signaling state space of the combinatory inhibitor experiment.

Figure S13: Proteins that contributed ≥2.5% to a PC in the combinatory inhibitor experiment.

Figure S14: Principal components PC1-3 visualized for DMSO controls (basal and activated) and 10 µM single inhibitor treatment.

Figure S15: Inhibition effect of 10 µM iSYK or 10 µM iNFκB in activated cells (combinatory inhibitor experiment).

Figure S16: Individual protein data exemplifying either strong repression across all inhibitor combinations that were investigated, or inhibitor-dependent differences in the strength of repression.

Figure S17: Phospho flow cytometry gating strategy.

Figure S18: Antibody barcode design for the ID-seq protocol.

Figure S19: Representative examples of successful antibody-DNA oligo conjugation for ID-seq.

Table S1: Antibody panel for ID-seq.

Tabel S2: Antibody panels for phospho-specific flow cytometry.

Table S3: Complete barcode and primer sequences for ID-seq.

Table S4: Well specific barcodes for ID-seq.

Table S5: Nextflex 8bp index barcodes for ID-seq.

## Notes

### Competing Interest Statement

The authors have declared no competing interest.

