## Supplementary Information for "Small molecule inhibitor combination treatment effectively represses global B-cell signaling in diffuse large B-cell lymphoma"

**Affiliations**

### Supplementary Materials

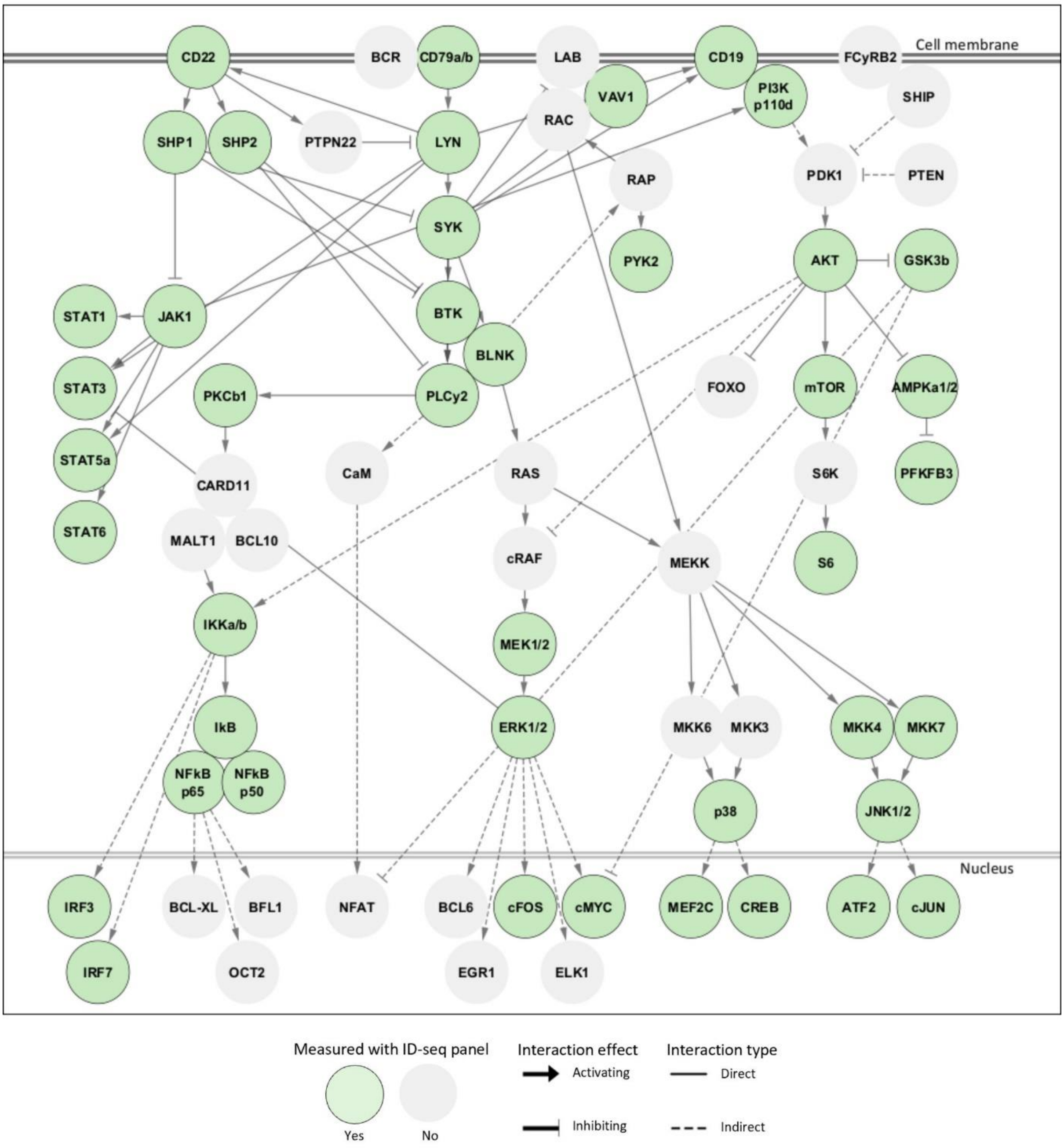


***Figure S1: Detailed schematic of the B-cell signaling network, with phospho-proteins measured in ID-seq panel indicated.***

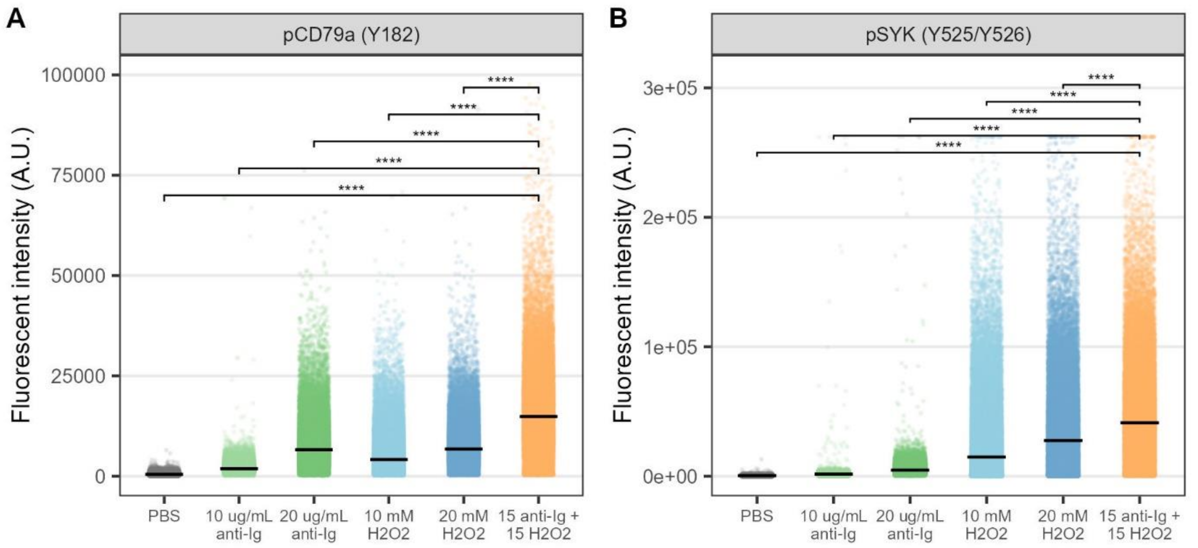


***Figure S2: Increased phosphorylation upon treatment with both anti-Ig and $H_2O_2$ combined compared to treatment with anti-Ig or $H_2O_2$ alone.*** *Significance was determined with the Kruskal-Wallis test ($p < 0.001$) and post-hoc Dunn's test with Benjamini-Hochberg (BH)-correction for multiple testing (comparing every condition with the 15 µg/mL anti-Ig + 15 mM $H_2O_2$ condition) (p-value threshold ≤ 0.05). A. CD79a. B. SYK. N > 5000 cells per condition. A.U. = arbitrary units.*

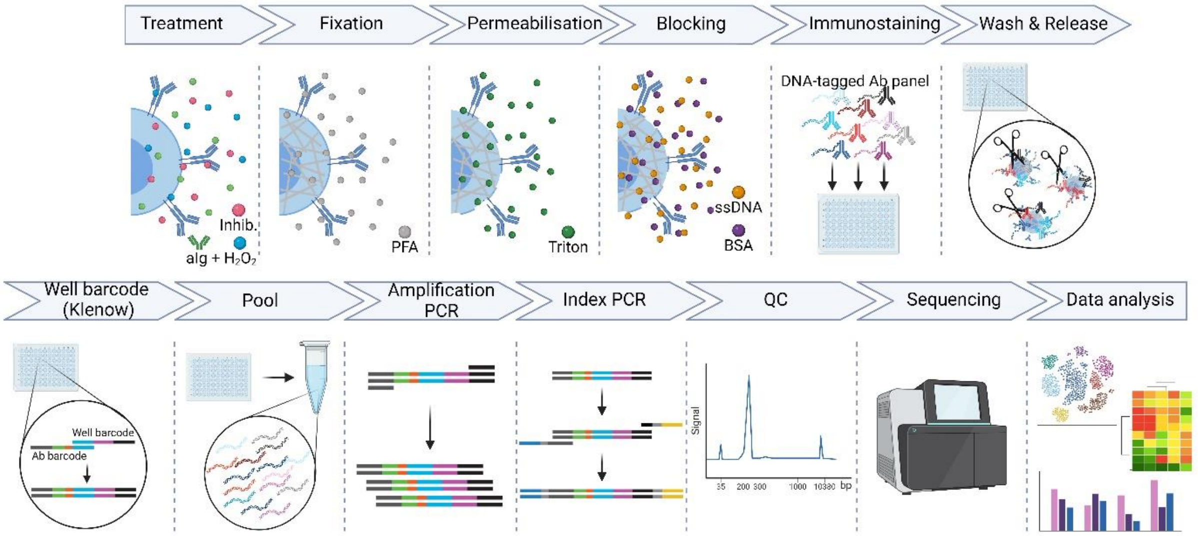


***Figure S3: Schematic overview of the experimental workflow of ID-seq.*** *Cells were treated with activating and inhibiting molecules. Then, cells were fixed with paraformaldehyde (PFA), and the membranes were permeabilized with Triton X-100. Blocking treatment with bovine serum albumin (BSA) and single stranded salmon sperm DNA (ssDNA) reduced unspecific binding. Then, immunostaining with a 111 DNA-labeled antibody panel was performed. The cells were washed, and the DNA barcode was released from the antibodies. A well-specific barcode was attached via a Klenow reaction. After pooling all samples and an amplification PCR, sequencing adapters were introduced in an index PCR step. The quality of the sequencing library was checked on a BioAnalyzer machine, and the sample was sequenced. Then, various data analysis approaches were applied, including (pre-)processing, differential expression analysis, and principal component analysis.*

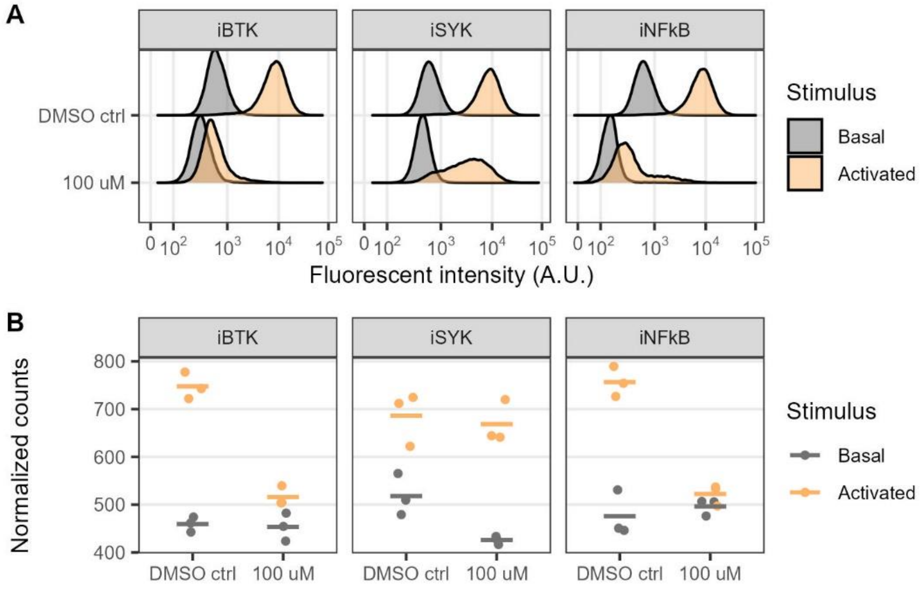


***Figure S4: Phospho flow cytometry measurements of CD79a (Y182) shows high correlation with ID-seq results.*** *After treatment and fixation, samples were split into two aliquots and subsequently measured with either phospho flow cytometry (A; N > 5000 cells per condition) or ID-seq (B; N = 3 biological replicates per condition).*

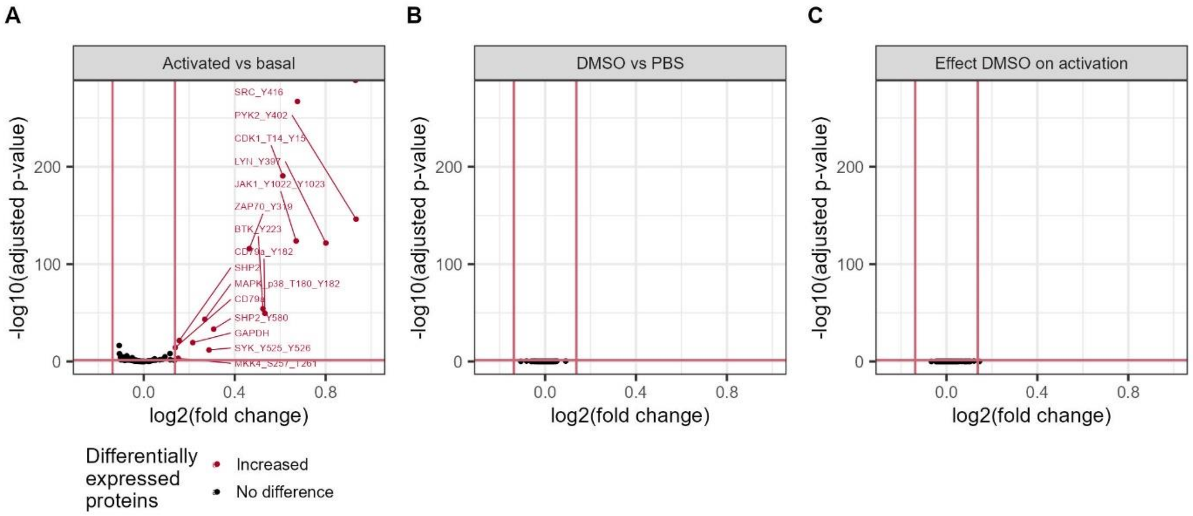


***Figure S5: No significant changes in (phospho-)protein levels upon treatment with DMSO compared to PBS.*** *N = 3 biological replicates per condition. Log2FC threshold ≥ log2(1.1), Wald test with BH-adjusted p-value threshold ≤ 0.05. A. Activated versus basal signaling. B. DMSO treatment versus PBS treatment. C. Effect of DMSO treatment on activated versus basal signaling.*

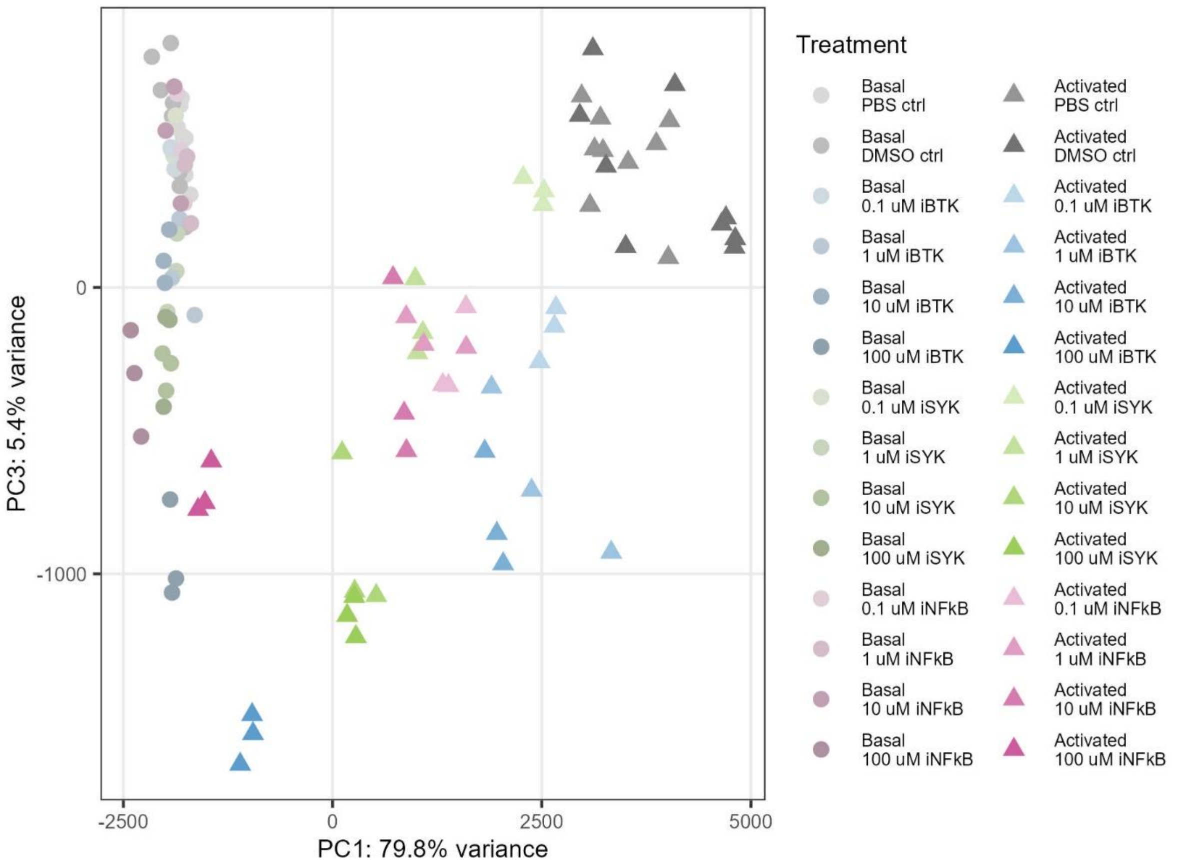


***Figure S6: Complete PCA signaling state landscape of the inhibitor dose experiment.*** *The PCA state landscape was determined with all samples in the experiment (N = 3 biological replicates per condition, except N = 9 biological replicates for DMSO controls). Subsequent visualization of conditions within this state landscape was performed with selected samples of interest.*

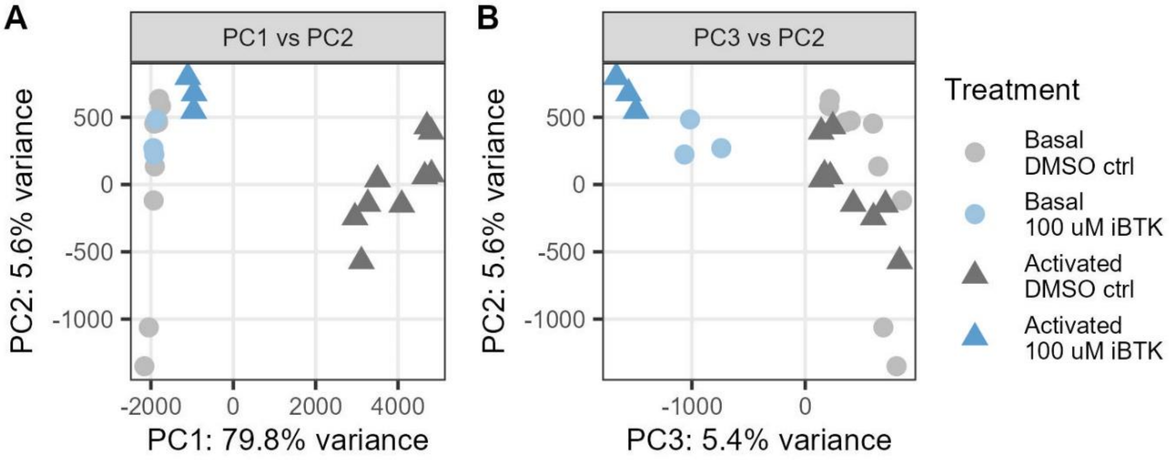


***Figure S7: Principal components PC1-3 visualized for DMSO controls (basal and activated) and 100 µM iBTK treatment.*** *N = 3 biological replicates per condition, except N = 9 biological replicates for DMSO controls. A. PC1 versus PC2. B. PC3 versus PC2.*

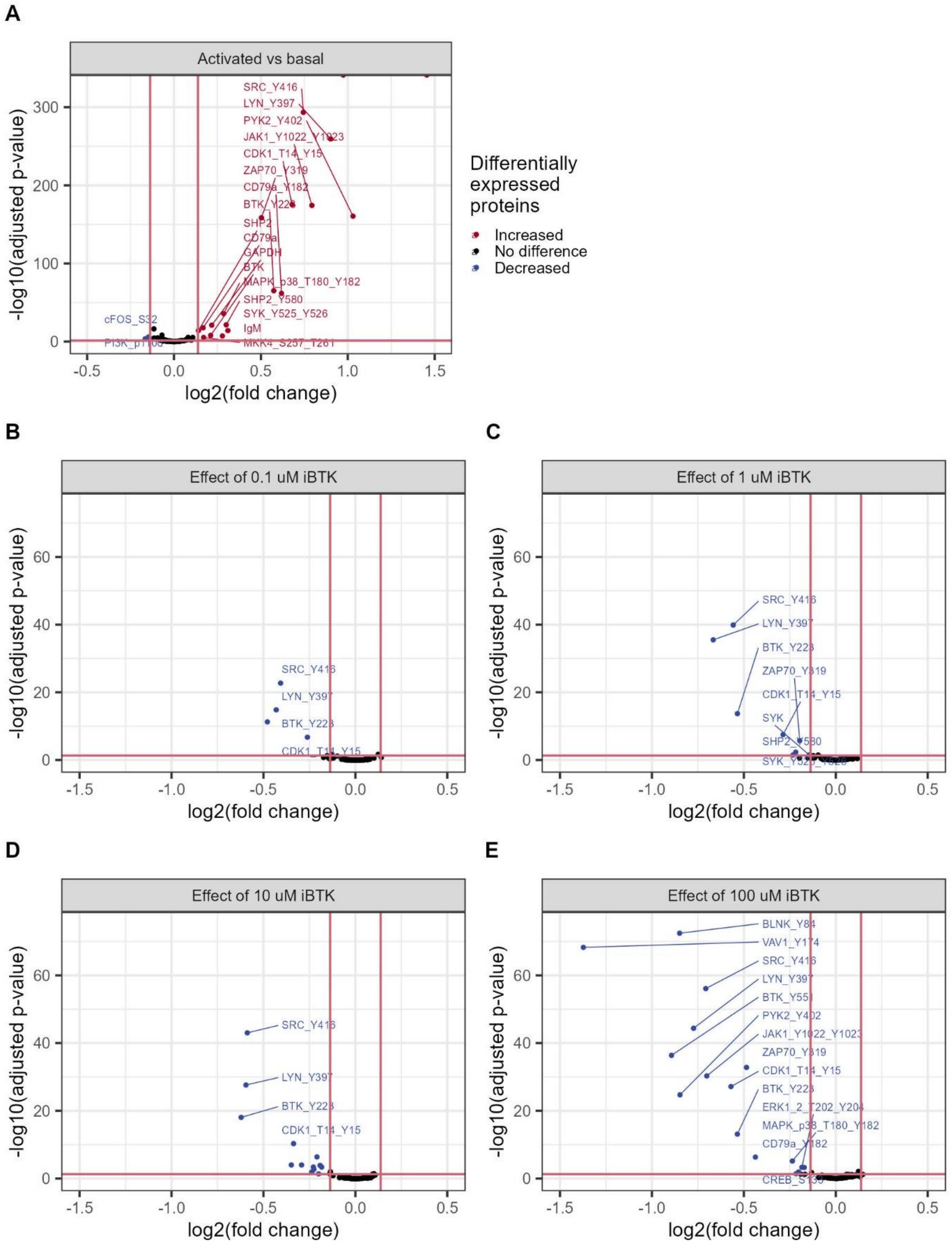


***Figure S8: Effect of iBTK dose on the comparison between activated and basal signaling.*** *Volcano plot visualization of the data represented in the heatmap of Fig. 3A. N = 3 biological replicates per condition. Log2FC threshold ≥ log2(1.1), Wald test with BH-adjusted p-value threshold ≤ 0.05.*

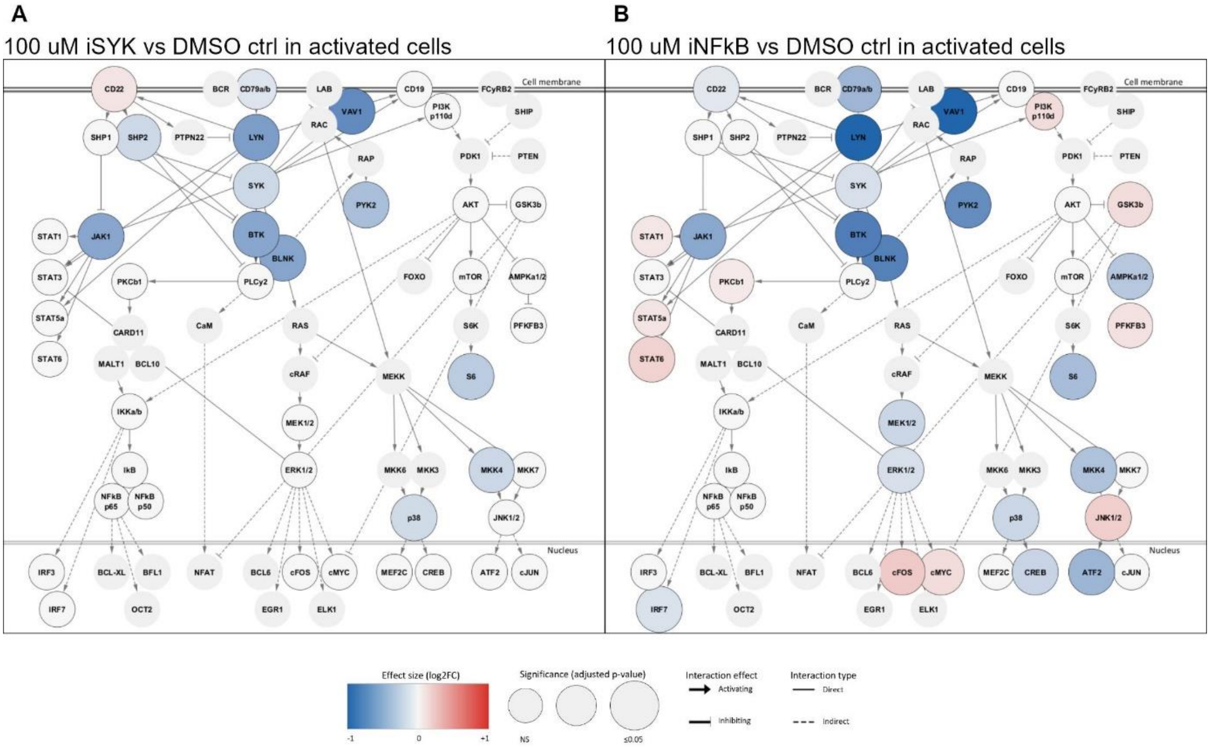


***Figure S9: Inhibition effect of 100 µM iSYK or 100 µM iNFκB versus DMSO control in activated cells.*** *Network visualization of DEA of 100 µM iSYK (A) or 100 µM iNFκB (B) treatment compared to DMSO control in activated cells. N = 3 biological replicates per condition. Wald test with BH-adjusted p-value threshold ≤ 0.05 (same threshold as Fig. 1D).*

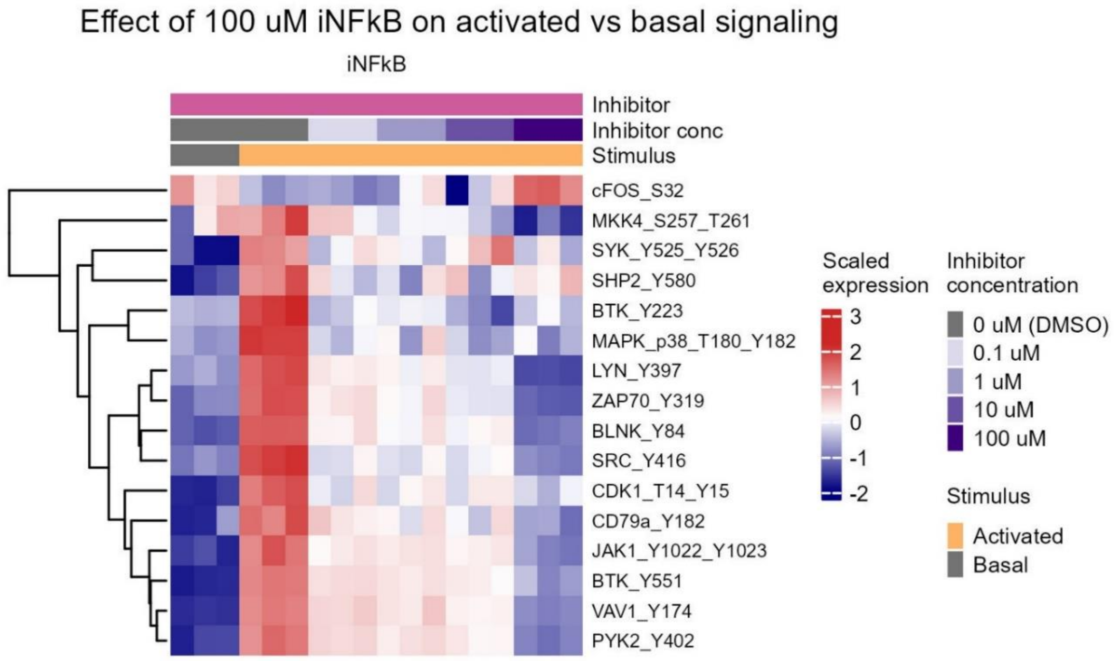


***Figure: S10: Effect of iNFκB dose on the comparison between activated and basal signaling.*** *Heatmap visualization of significant proteins from the DEA of activated versus basal treatment and the effect of iNFκB dose on the activated versus basal comparison. N = 3 biological replicates per condition. Log2FC threshold ≥ log2(1.1), Wald test with BH-adjusted p-value threshold ≤ 0.05.*

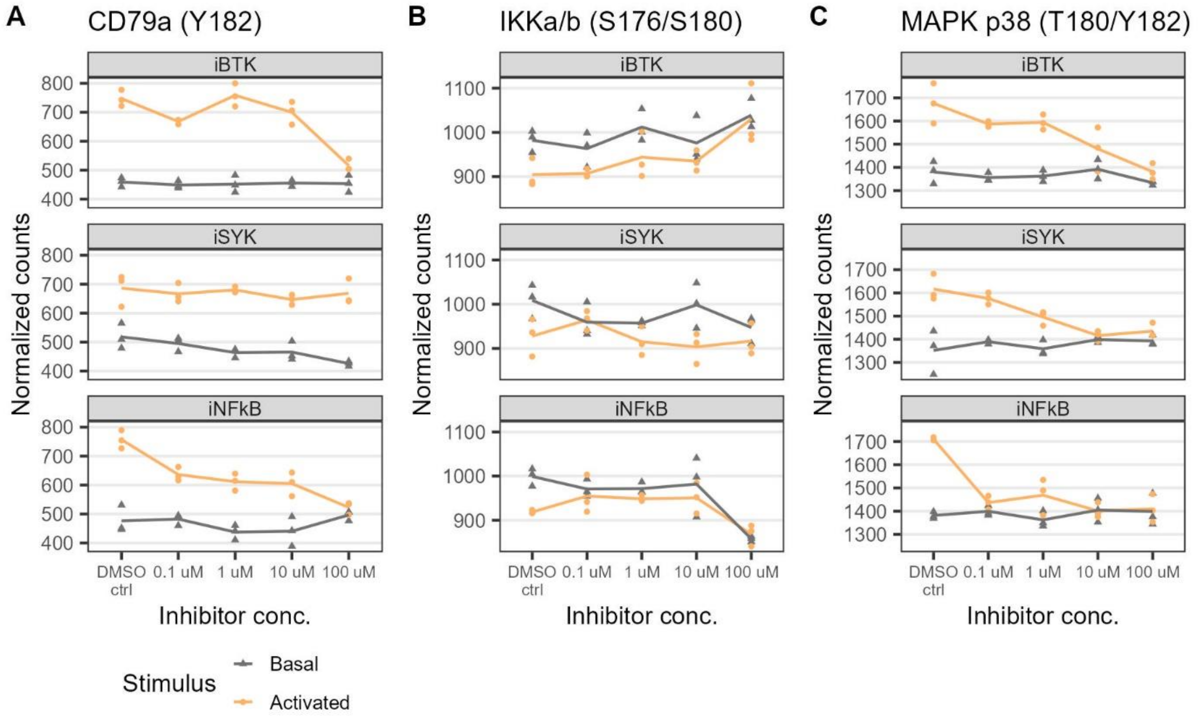


***Figure S11: Different dose-response patterns in phosphorylation levels of CD79a, IKKa/b, and MAPK p38 upon activation and inhibition by iBTK, iSYK, and iNFκB.*** *N = 3 biological replicates per condition (individual replicates and mean normalized ID-seq count data). Significance was determined in DEA (see Fig. 3A, Fig. 4B, and Suppl. Fig. S10). A. CD79a. B. IKKa/b. C. MAPK p38.*

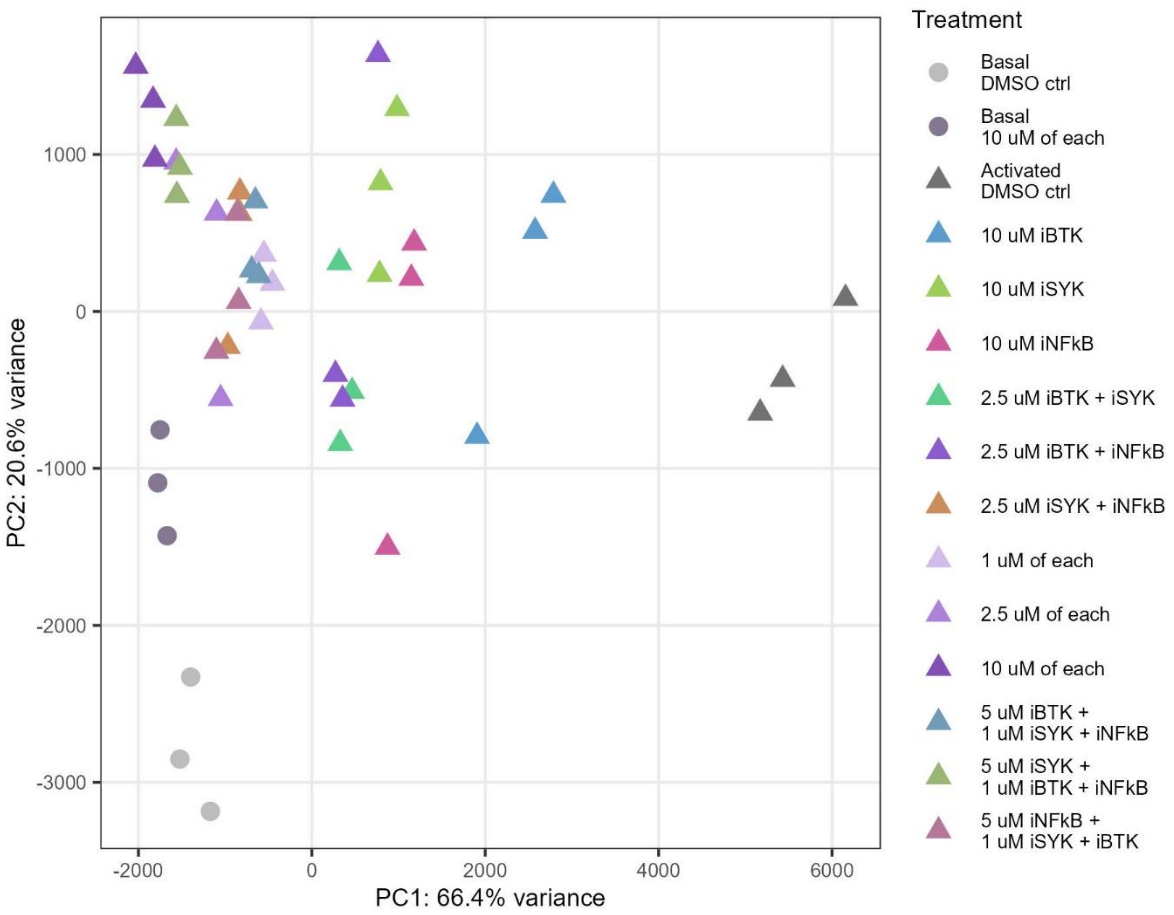


***Figure S12: Complete PCA signaling state landscape of the combinatory inhibitor experiment.*** *The PCA state landscape was determined with all sample in the experiment. N = 3 biological replicates per condition. Subsequent visualization of conditions within this signaling state landscape was performed with selected samples of interest.*

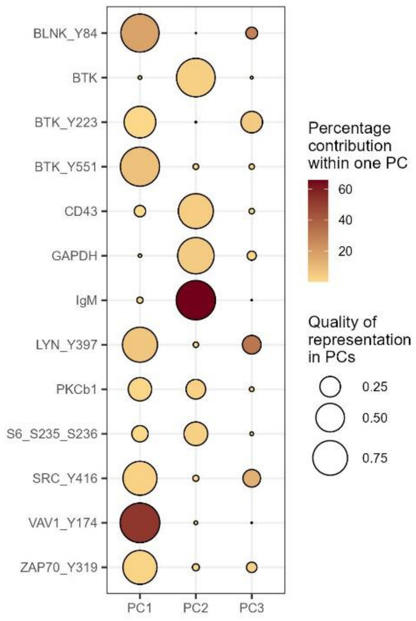


***Figure S13: Proteins that contributed ≥2.5% to a PC in the combinatory inhibitor experiment.***

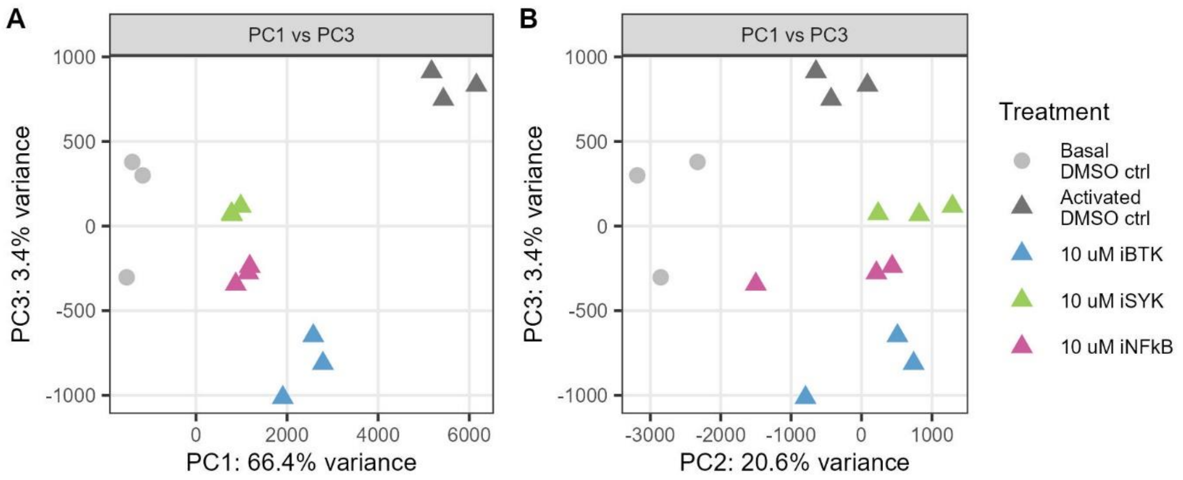


***Figure S14: Principal components PC1-3 visualized for DMSO controls (basal and activated) and 10 µM single inhibitor treatment.*** *N = 3 biological replicates per condition. A. PC1 versus PC3. B. PC2 versus PC3.*

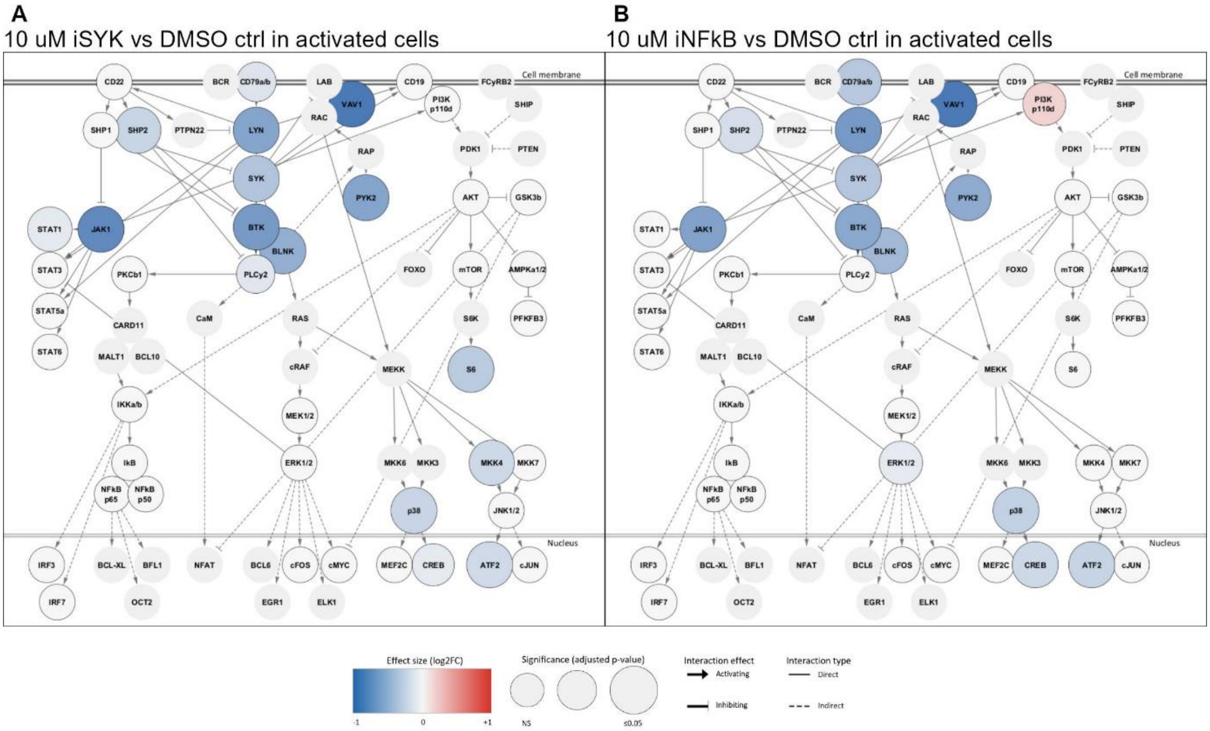


***Figure S15: Inhibition effect of 10 µM iSYK or 10 µM iNFκB in activated cells (combinatory inhibitor experiment).*** *Network visualization of the DEA of 10 µM iSYK (A) or 10 µM iNFκB (B) compared to DMSO control in activated cells. N = 3 biological replicates per condition. Wald test with BH-adjusted p-value threshold ≤ 0.05.*

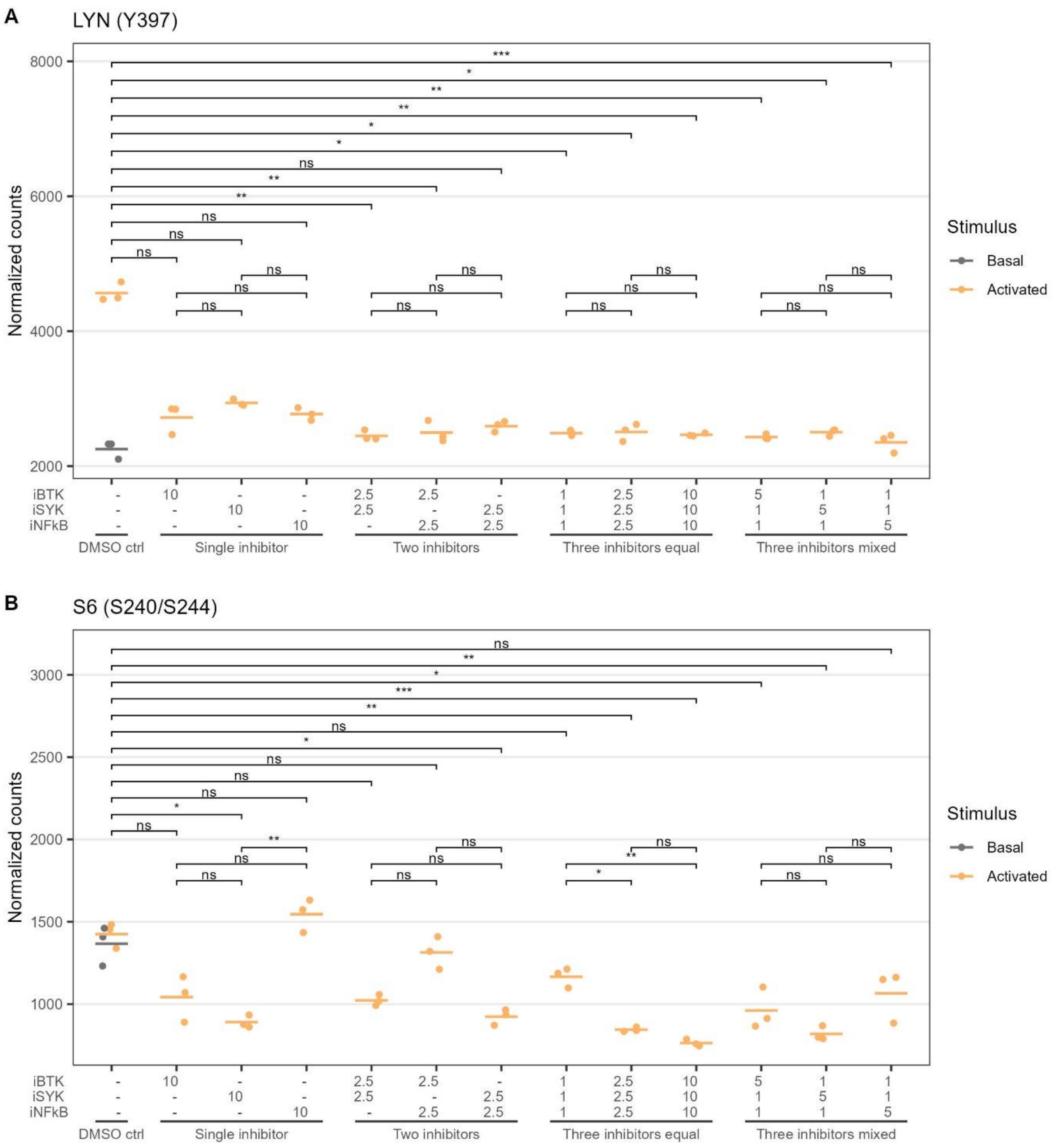


***Figure S16: Individual protein data exemplifying either strong repression across all inhibitor combinations that were investigated, or inhibitor-dependent differences in the strength of repression.*** *Phosphorylation levels of LYN (A) and S6-S240/S244 (B) (N = 3 biological replicates and mean normalized ID-seq count data). Significance was determined with the Kruskal-Wallis test ($p < 0.001$) and post-hoc Dunn's test with BH-correction for multiple testing (comparing every condition with the DMSO control condition and each condition within a treatment group) p-value threshold ≤ 0.05).*

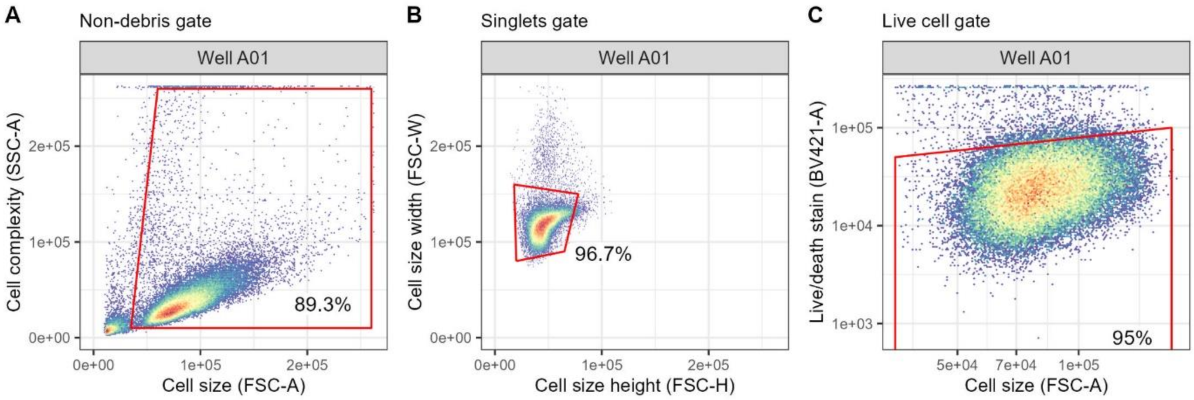


***Figure S17: Phospho flow cytometry gating strategy.*** *A. Non-debris gate with cell size (forward scatter [FSC]) and cell complexity (side scatter [SSC]). B. Singlets gate with cell size height (FSC-H) and cell size width (FSC-W). C. Live cell gate with cell size (FSC) and live/death stain (cleaved Caspase 3 + cleaved PARP [BV421]).*

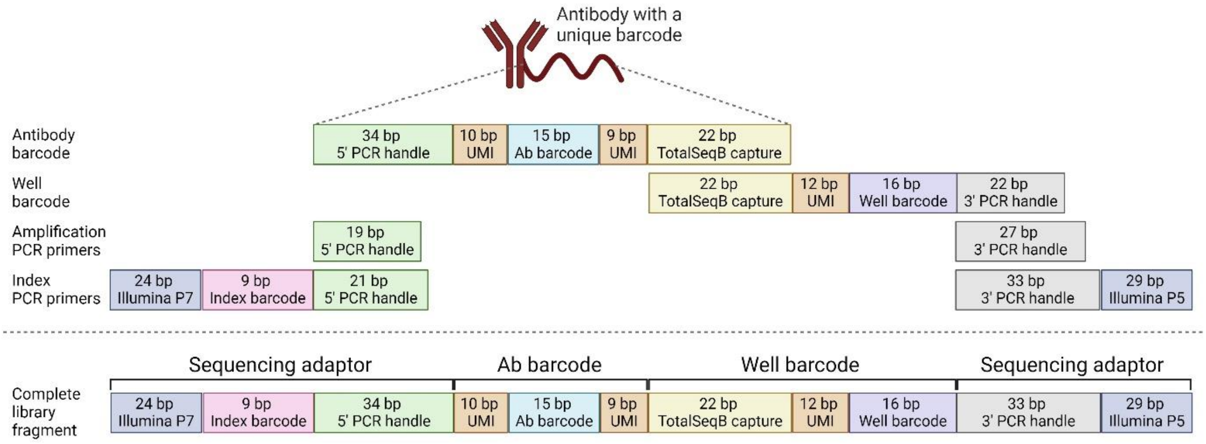


***Figure S18: Antibody barcode, well barcode, and primer designs for the ID-seq protocol.*** *The antibody barcode oligo consists of a 5' PCR handle, unique molecular identifier (UMI), antibody-specific barcode, UMI, and TotalSeqB capture sequence. The well barcode consists of the TotalSeqB capture sequence, UMI, well-specific barcode, and a 3' PCR handle. The amplification PCR primers cover the 5' and 3' PCR handles. The index PCR primers contain, in addition to the 5' and 3' PCR handles, also the Illumina P7 and P5 sequencing adaptors, and one Nextera index barcode. See Suppl. Tables S1, S3, S4, and S5 for all barcode and primer sequences.*

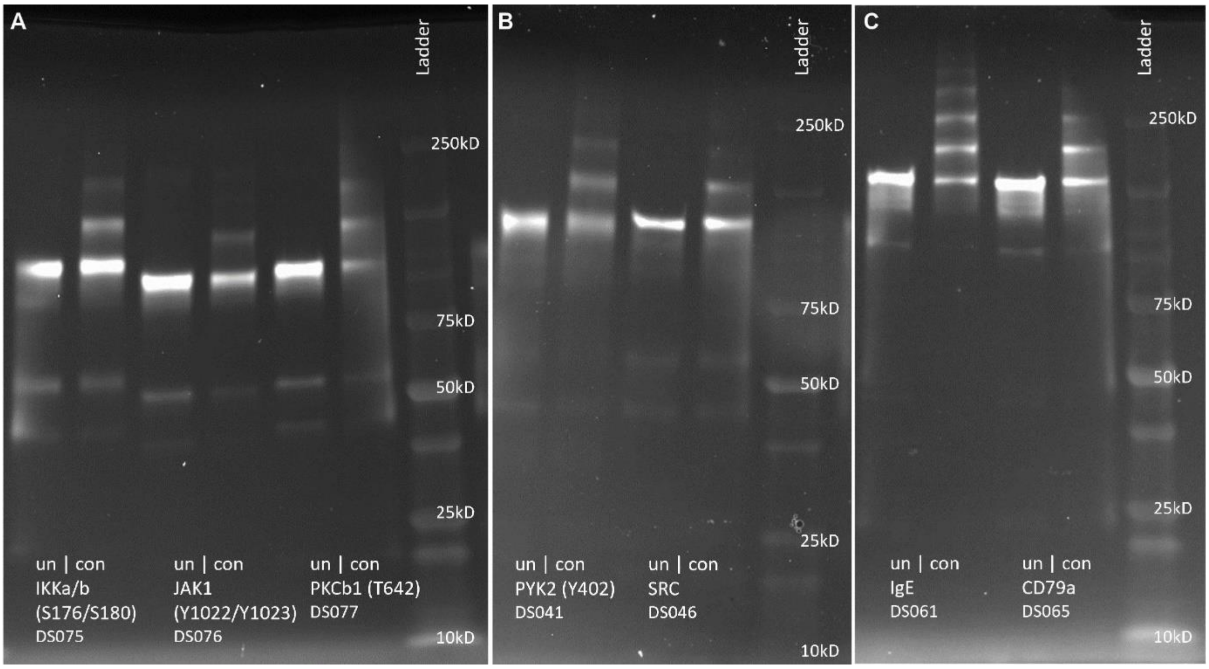


***Figure S19: Representative examples of successful antibody-DNA oligo conjugation for ID-seq.*** *Samples were run on a non-reducing SDS-PAGE gel. Un = unconjugated antibody, con = conjugated antibody, DS number refers to Suppl. Table S1. A-C Three different gels from different conjugation batches.*

***Table S1: Antibody panel for ID-seq.*** *NA = not applicable.*

| Ab nr. | Target | Ab barcode sequence | Source | Catalog number | Clone/Identifier |
|---|---|---|---|---|---|
| DS001 | AKT | TTACATCGAGAATCA | Cell Signaling Technologies | 4685 | Clone:11EE7<br>RRID:AB_2225340 |
| DS002 | AKT (S473) | TACAGGTACACATTG | Cell Signaling Technologies | 4060 | Clone:D9E<br>RRID:AB_2315049 |
| DS003 | AKT (T308) | CAGCTGATCTAATTC | Cell Signaling Technologies | 13038 | Clone:D25E6<br>RRID:AB_2629447 |
| DS068 | AMPKa1/2 (T183/T172) | GCGCTGTGCCGAGGA | Thermo Fisher Scientific | 701068 | Clone:10H2L20<br>RRID:AB_2532369 |
| DS069 | AMPKb1 (S182) | TAGTATCTATAGATC | Thermo Fisher Scientific | 700241 | Clone:9H26L42<br>RRID:AB_2532303 |
| DS081 | ATF2 (T71) | TCCGATCCTGGTCCG | Abcam | ab242381 | Clone:E268<br>RRID:AB_725567 |
| DS004 | BAD (S112) | AGGTTATATGTCGAG | Cell Signaling Technologies | 5284 | Clone:40A9<br>RRID:AB_560884 |
| DS005 | BAD (S136) | GCCTAACATGCTTCT | Cell Signaling Technologies | 4366 | Clone:D25H8<br>RRID:AB_10547878 |
| DS006 | BIM | CACATATAGAATTAG | Cell Signaling Technologies | 2933 | Clone:C34C5<br>RRID:AB_1030947 |
| DS007 | BLNK | AGTAATGCATACCTG | Cell Signaling Technologies | 36438 | Clone:D3P2H<br>RRID:AB_2799101 |
| DS063 | BLNK (Y84) | ACTAGGTTAACCTCT | BD Biosciences | 558366 | Clone:J117-1278<br>RRID:AB_647263 |
| DS008 | BTK | TACCAACTGGTCTGA | Cell Signaling Technologies | 94988 | Clone:D3H5<br>RRID:AB_10950506 |
| DS054 | BTK (Y223) | GACGGCCTCGGCAAG | Biolegend | 601702 | Clone:A16128B<br>RRID:AB_2715907 |
| DS009 | BTK (Y551) | GCCGTCCGCGCGTAG | Cell Signaling Technologies | 18805 | Clone:E5Y6N<br>RRID:AB_3095069 |
| DS010 | Caspase 3 (D175) | GTGCAAGAGTTGGCG | Cell Signaling Technologies | 94530 | Clone:5A1E<br>RRID:AB_3076239 |
| DS082 | CD10 | CTTGCGAGTAGGATT | Abcam | ab243934 | Clone:SP179<br>RRID: NA |
| DS083 | CD138 | ACCATAGACGATCAT | Abcam | ab226108 | Clone:EPR6454<br>RRID:AB_2889392 |
| DS084 | CD19 | AGGACAACACTCAGT | Abcam | ab271904 | Clone:EPR5906 |

| | | | | | |
|---|---|---|---|---|---|
| | | | | | RRID:AB_2801636 |
| DS064 | CD20 | CCTGAGTGAGGATAG | BD Biosciences | 555677 | Clone:H1<br>RRID:AB_396030 |
| DS091 | CD22 (Y822) | TCAAGCCTCAGGCAG | R&D Systems | MAB7290 | Clone:726830<br>RRID:AB_11128833 |
| DS092 | CD24 | GAGCATAAGTGGTTA | R&D Systems | MAB5247-100 | Clone:ML5<br>RRID: NA |
| DS085 | CD27 | CCGCATGAGGCCTGC | Abcam | ab256583 | Clone:EPR8569<br>RRID:AB_11155136 |
| DS093 | CD37 | GACGAACGTCCGCGA | R&D Systems | MAB4625 | Clone:424925<br>RRID:AB_2228783 |
| DS055 | CD38 | GATTGGCTACTCAAT | Biolegend | 303535 | Clone:HIT2<br>RRID:AB_2562819 |
| DS086 | CD43 | GTTAAGTCATACATA | Abcam | ab238799 | Clone:SP55<br>RRID:AB_10710687 |
| DS065 | CD79a | AACCTACTCTCAAGG | BD Biosciences | 555934 | Clone:HM47<br>RRID:AB_396231 |
| DS011 | CD79a (Y182) | CTAATGTCACAAGTA | Cell Signaling Technologies | 14732 | Clone:D1B9<br>RRID:AB_2798591 |
| DS012 | CDC2 (Y15) | GAAGAAGCGTTATTC | Cell Signaling Technologies | 4539 | Clone:10A11<br>RRID:AB_560953 |
| DS070 | CDK1 (T14/Y15) | CATTGTGACTTCACC | Thermo Fisher Scientific | 701808 | Clone:17H29L7<br>RRID:AB_2609690 |
| DS071 | CDK4 (T172) | GTTCAAGCTTAGATA | Thermo Fisher Scientific | 702556 | Clone:9H2L7<br>RRID:AB_2632989 |
| DS072 | CDK6 (T177) | TGCATAGCCTGTGGA | Thermo Fisher Scientific | 711588 | Clone:16HCLC<br>RRID:AB_2632988 |
| DS013 | c-FOS (S32) | TGGTGACAAGTATCT | Cell Signaling Technologies | 5348 | Clone:D82C12<br>RRID:AB_10557109 |
| DS014 | c-JUN (S63) | GCCAACGGCTATATG | Cell Signaling Technologies | 91952 | Clone:E6I7P<br>RRID:AB_2893112 |
| DS073 | c-MYC | TAGAATCGCTACAAC | Thermo Fisher Scientific | 700648 | Clone:27H46L35<br>RRID:AB_2532334 |
| DS015 | c-MYC (T58) | CGAGGTACATCTTGT | Cell Signaling Technologies | 46650 | Clone:E4Z2K<br>RRID: NA |
| DS016 | CREB (S133) | TCTCTATGAATGTTG | Cell Signaling Technologies | 9198 | Clone:87G3<br>RRID:AB_2561044 |

| | | | | | |
|---|---|---|---|---|---|
| DS056 | Cyclin A2 | TGAATGCAAGTAATT | Biolegend | 644001 | Clone:E23.1<br>RRID:AB_2071969 |
| DS057 | Cyclin B1 | CACGATCCTCGTTAC | Biolegend | 647902 | Clone:V152<br>RRID:AB_2244210 |
| DS017 | Cyclin D1 | TAACGTCACAACATA | Cell Signaling Technologies | 66467 | Clone:E3P5S<br>RRID:AB_2827374 |
| DS074 | Cyclin E | AATGTTACGTGACCT | Thermo Fisher Scientific | 321600 | Clone:HE12<br>RRID:AB_2533067 |
| DS108 | ELK1 | TCCTGAGTGTCAATA | Abcam | ab227114 | Clone:E277<br>RRID: NA |
| DS018 | ERK1/2 | AGATCTATTGGCAGT | Cell Signaling Technologies | 4695 | Clone:137F5<br>RRID:AB_390779 |
| DS019 | ERK1/2 (T202/Y204) | TTAGGTGTACACGTT | Cell Signaling Technologies | 4370 | Clone:D13.14.4E<br>RRID:AB_2315112 |
| DS113 | FOXO1a/3a (T24/T32) | AACTGTCGCCACGTC | Abcam | ab312327 | Clone:EPR28359-78<br>RRID: NA |
| DS058 | GAPDH | ACAGAGAGTGTGACG | Biolegend | 607902 | Clone:W17079A<br>RRID:AB_2734503 |
| DS020 | GSK-3b (S9) | TAGGAGGTATCCTCA | Cell Signaling Technologies | 5558 | Clone:D85E12<br>RRID:AB_10013750 |
| DS059 | Histone H2A.X (S139) | TGACCTCCATAATAA | Biolegend | 613402 | Clone:2F3<br>RRID:AB_315795 |
| DS021 | Histone H3 | GAGTTACCATTGAAC | Cell Signaling Technologies | 60932 | Clone:D1H2<br>RRID:AB_10544537 |
| DS060 | Histone H3 (S10) | GTGGCTGAGCTGACG | Biolegend | 650802 | Clone:11D8<br>RRID:AB_10896911 |
| DS087 | IgA | AGGCGTGCACGTGGT | Abcam | ab214003 | Clone:EPR5367-76<br>RRID: NA |
| DS066 | IgD | AATTCAGGCGACGTG | BD Biosciences | 555776 | Clone:IA6-2<br>RRID:AB_396111 |
| DS061 | IgE | TTACTACGTAATACC | Biolegend | 325502 | Clone:MHE-18<br>RRID:AB_830847 |
| DS067 | IgG | ATAGGTATATCCATG | BD Biosciences | 555784 | Clone:G18-145 |
| DS062 | IgM | TTCGGAACGCGCACA | Biolegend | 314502 | Clone:MHM-88<br>RRID:AB_493003 |
| DS022 | IkB | CATAGCCAGGTTATC | Cell Signaling Technologies | 4814 | Clone:L35A5<br>RRID:AB_390781 |

| | | | | | |
|---|---|---|---|---|---|
| DS075 | IKKa/b (S176/S180) | TTCATCTCCAGTACG | Thermo Fisher Scientific | 701643 | Clone:7H17L17<br>RRID:AB_2532498 |
| DS023 | IRF3 (S396) | AGCAGGCTTGAAGGA | Cell Signaling Technologies | 29047 | Clone:D6O1M<br>RRID:AB_2773013 |
| DS107 | IRF7 (S477/S479) | GGCAGCCTTCTATTG | BD Biosciences | 558621 | Clone:K47-671<br>RRID:AB_396119 |
| DS094 | Isotype Mouse IgG1 | GTCACACTTAAGACA | R&D Systems | MAB002R | Clone:11711R<br>RRID: NA |
| DS024 | Isotype Rabbit IgG | CTGGTCGAACTTCGT | Cell Signaling Technologies | 3900 | Clone:DA1E<br>RRID:AB_1550038 |
| DS095 | JAK1 | CTCGTCGCGCAATGC | R&D Systems | MAB4260 | Clone:413104<br>RRID:AB_2128403 |
| DS076 | JAK1 (Y1022/Y1023) | CGGCTCACCGCGTCT | Thermo Fisher Scientific | 700028 | Clone:59H4L5<br>RRID:AB_2532272 |
| DS025 | JNK1/2 (T183/Y185) | ACAGGCCGCATACTT | Cell Signaling Technologies | 9255 | Clone:G9<br>RRID:AB_2307321 |
| DS026 | Ki-67 | AGAGCTAGGATCGGA | Cell Signaling Technologies | 9129 | Clone:D3B5<br>RRID:AB_2687446 |
| DS096 | LYN | CGCGGCCGGTTAGGA | R&D Systems | MAB3206 | Clone:338716<br>RRID:AB_2138374 |
| DS027 | LYN (Y397) | ATAATCATTACGTGG | Cell Signaling Technologies | 70926 | Clone:E5L3D<br>RRID:AB_2924371 |
| DS028 | MAPK p38 (T180/Y182) | TTGAAGGTAATACAG | Cell Signaling Technologies | 4511 | Clone:D3F9<br>RRID:AB_2139682 |
| DS029 | MEF2C | TATCCGATTCGCGCG | Cell Signaling Technologies | 5030 | Clone:D80C1<br>RRID:AB_10548759 |
| DS030 | MEK1/2 | ATTACCGATATACAA | Cell Signaling Technologies | 8727 | Clone:D1A5<br>RRID:AB_10829473 |
| DS097 | MEK1/2 (S218/S222) | AGCTTCACGGAATCC | R&D Systems | MAB8407 | Clone:1020E<br>RRID: NA |
| DS031 | MKK4 (S257/T261) | CGAGTCCAGGCCGGA | Cell Signaling Technologies | 4514 | Clone:C36C11<br>RRID:AB_2140946 |
| DS032 | MKK7 (S271/T275) | CAAGCGCGGCTTCCG | Cell Signaling Technologies | 4171 | Clone:polyclonal<br>RRID:AB_2250408 |
| DS033 | mTOR (S2448) | GCTAGCGTAATGTGG | Cell Signaling Technologies | 5536 | Clone:D9C2<br>RRID:AB_10691552 |
| DS034 | NFκB p50 | CATGGAGGCGAGGTT | Cell Signaling Technologies | 13586 | Clone:D4P4D |

| | | | | | |
|---|---|---|---|---|---|
| | | | | | RRID:AB_2665516 |
| DS035 | NFκB p65 | TGACACCTGAATATT | Cell Signaling Technologies | 69994 | Clone:D14E12<br>RRID:AB_10859369 |
| DS036 | NFκB p65 (S536) | ACGAAGCGTGGCAGA | Cell Signaling Technologies | 3033 | Clone:93H1<br>RRID:AB_331284 |
| DS111 | NFκB RelB (S552) | GAGTGAGGCTCAGCT | Cell Signaling Technologies | 17616 | Clone:D41B9<br>RRID:AB_10622001 |
| DS109 | p53 (S15) | TAACGGACTGACACG | Cell Signaling Technologies | 56979 | Clone:16G8<br>RRID:AB_331741 |
| DS037 | p90RSK1 (T359) | TAAGGACGCGTACGG | Cell Signaling Technologies | 8753 | Clone:D1E9<br>RRID:AB_2783561 |
| DS038 | PARP (D214) | ACCAGTTACAGGACT | Cell Signaling Technologies | 5625 | Clone:D64E10<br>RRID:AB_10699459 |
| DS088 | PFKFB3 (S461) | CGCTGCGGCGAGTTG | Abcam | ab232498 | Clone:EPR19735<br>RRID: NA |
| DS039 | PI3K p110d | CCAATCGATATGAGC | Cell Signaling Technologies | 19591 | Clone:E2T2N<br>RRID: NA |
| DS089 | PKC-b1 | ACCTTCTGTTCTGCC | Abcam | ab232518 | Clone:EPR18512<br>RRID: NA |
| DS077 | PKC-b1 (T642) | CCGAACGCAGCCGTA | Thermo Fisher Scientific | 702430 | Clone:3H8L1<br>RRID:AB_2662390 |
| DS040 | PLCg1 (Y783) | ATGCCGAATGAGTCG | Cell Signaling Technologies | 14008 | Clone:D6M9S<br>RRID:AB_2728690 |
| DS098 | PLCg2 | TCTATTGGATAGGAT | R&D Systems | MAB3716 | Clone:346404<br>RRID:AB_2163529 |
| DS099 | PLCg2 (Y753) | CCATCAGGACGGAGT | R&D Systems | MAB37161 | Clone:790623<br>RRID: NA |
| DS100 | PLCg2 (Y759) | TGGTTGATCGTATCT | R&D Systems | MAB7377 | Clone:744757<br>RRID: NA |
| DS041 | PYK2 (Y402) | TTAGTGGTCCGGTTA | Cell Signaling Technologies | 3291 | Clone:polyclonal<br>RRID:AB_2300530 |
| DS042 | Rb (S807/S811) | TAGACCATGCCGCCT | Cell Signaling Technologies | 8516 | Clone:D20B12<br>RRID:AB_11178658 |
| DS043 | S6 (S235/S236) | GACGGCCGCGCCGTC | Cell Signaling Technologies | 4858 | Clone:D57.2.2E<br>RRID:AB_916156 |
| DS044 | S6 (S240/S244) | AATGCACCGAAGTGC | Cell Signaling Technologies | 5364 | Clone:D68F8<br>RRID:AB_10694233 |

| | | | | | |
|---|---|---|---|---|---|
| DS101 | SHP1 | TCTCGCACCACGCGG | R&D Systems | MAB1878 | Clone:255402<br>RRID:AB_2251548 |
| DS102 | SHP2 | TGTCAGCGACTGCGA | R&D Systems | MAB1894 | Clone:255509<br>RRID:AB_2175228 |
| DS045 | SHP2 (Y580) | GGTCATAGGAAGTGT | Cell Signaling Technologies | 5431 | Clone:D66F10<br>RRID:AB_10693803 |
| DS046 | SRC | ACGTGAGGAGGCAGT | Cell Signaling Technologies | 2109 | Clone:36D10<br>RRID:AB_2106059 |
| DS103 | SRC (Y416) | GCAACTGGCATCATC | R&D Systems | MAB2685 | Clone:1246F<br>RRID: NA |
| DS047 | STAT1 | GCGGCTCTGTTGATG | Cell Signaling Technologies | 65748 | Clone:D1K9Y<br>RRID:AB_2737027 |
| DS078 | STAT1 (Y701) | GACGGCACCAAGTTC | Thermo Fisher Scientific | 333400 | Clone:ST1P-11A5<br>RRID:AB_2533113 |
| DS048 | STAT3 | TTGTCACGGTAATAA | Cell Signaling Technologies | 12640 | Clone:D3Z2G<br>RRID:AB_2629499 |
| DS104 | STAT3 (S727) | TTATGGAGTGTAACA | R&D Systems | MAB4934 | Clone:788335<br>RRID: NA |
| DS049 | STAT3 (Y705) | ATCGAACCGACAGAG | Cell Signaling Technologies | 9145 | Clone:D3A7<br>RRID:AB_2491009 |
| DS105 | STAT5a | CCTAGTTACCGAGCA | R&D Systems | MAB21741 | Clone:251610<br>RRID:AB_2196766 |
| DS079 | STAT5a (Y694) | ATAGCGGCTCTATCG | Thermo Fisher Scientific | 701063 | Clone:6H5L15<br>RRID:AB_2532365 |
| DS106 | STAT6 | CAGCCTACAATATGC | R&D Systems | MAB2167 | Clone:253906<br>RRID:AB_2271214 |
| DS080 | STAT6 (Y641) | AGTACAGGTGACTTA | Thermo Fisher Scientific | 700247 | Clone:46H1L12<br>RRID:AB_10562728 |
| DS050 | SYK | GGTCGACTAGGTCGG | Cell Signaling Technologies | 13198 | Clone:D3Z1E<br>RRID:AB_2687924 |
| DS051 | SYK (Y525/Y526) | GTACTGACTATGCTT | Cell Signaling Technologies | 2710 | Clone:C87C1<br>RRID:AB_2197222 |
| DS052 | TAK1 (T184/T187) | CGTGCCAGCGAATTC | Cell Signaling Technologies | 4508 | Clone:90C7<br>RRID:AB_561317 |
| DS090 | VAV1 (Y174) | GCAAGCAATGCACGC | Abcam | ab238424 | Clone:EP510Y<br>RRID:AB_1524546 |
| DS053 | ZAP70 (Y319) | CTCAAGCATTATCAT | Cell Signaling Technologies | 2717 | Clone:65E4 |

| | | | | | |
|---|---|---|---|---|---|
| | | | | | RRID:AB_2218658 |

***Table S2: Antibody panels for phospho-specific flow cytometry.***

| Target | Fluorophore | Dilution | Source | Catalog number | Clone/Identifier |
|---|---|---|---|---|---|
| Active Caspase 3 | V450 | 1/500 | BD Biosciences | 560627 | Clone:C92-605<br>RRID:AB_1727415 |
| Cleaved PARP (D214) | BV421 | 1/500 | BD Biosciences | 564129 | Clone:F21-852<br>RRID:AB_2738611 |
| pSYK (Y525/Y526) | PE | 1/200 | Cell Signalling Technologies | 6485 | Clone:C87C1<br>RRID:AB_11220429 |
| pPLCγ2 (Y759) | Alexa 647 | 1/100 | BD Biosciences | 558498 | Clone:K86-689.37<br>RRID:AB_647139 |
| pCD79a (Y182) | Alexa 488 | 1/200 | Cell Signalling Technologies | 52821 | Clone:D1B9<br>RRID:AB_2799422 |

***Table S3: Complete barcode and primer sequences for ID-seq.*** *Green = 5' PCR handle. Orange = unique molecular identifier (UMI) sequence. Light blue = Antibody specific barcode (see Suppl. Table S2). Yellow = TotalSeqB capture sequence. Grey = 3' PCR handle. Purple = Well specific barcode (see Suppl. Table S4). Red = Nextflex 8bp index barcode (see Suppl. Table S5).*

| **Barcode/ Primer** | **Sequence 5' > 3'** | **Length (bp)** |
|---|---|---|
| Antibody barcode complete | GTGACTGGAGTTCAGACGTGTGCTCTTCCGATCT[10bp UMI][15bp Ab barcode][9bp UMI]GCTTTAAGGCCGGTCCTAGCAA | 90 |
| Well barcode complete | GTCAGATGTGTATAAGAGACAG[16bp well barcode][12bp UMI]TTGCTAGGACCGGCCTTAAAGC | 72 |
| Amplification PCR forward | GCAGCGTCAGATGTGTATAAGAGACAG | 27 |
| Amplification PCR reverse | GTGACTGGAGTTCAGACGT | 19 |
| Index PCR forward | AATGATACGGCGACCACCGAGATCTACACTCGTCGGCAGCGTCAGATGTGTATAAGAGACAG | 62 |
| Index PCR reverse | CAAGCAGAAGACGGCATACGAGAT[8bp index barcode]GTGACTGGAGTTCAGACGTGT | 53 |

*Table S4: Well specific barcodes for ID-seq.*

| Barcode | Sequence 5' > 3' |
|---|---|
| A01 | AATAACACCTCGAATC |
| A02 | TCGTTATGTCCTGCTT |
| A03 | ACATCCCAGGTAGCAC |
| A04 | AGGATAACAGTCATGT |
| A05 | ATTCCATGTTAGGTAA |
| A06 | CATTGCCTCTAAACTG |
| A07 | TAAGCTCTCCCTCAGT |
| A08 | GAACGTTCAGGTGGAT |
| A09 | GCCATTCGTTACATCG |
| A10 | GTAGAGGTCGTTACCC |
| A11 | TACCCACAGGGCATGT |
| A12 | TCCTCCCCAGGCGTTC |
| B01 | ACTATCACTTGGCATG |
| B02 | TGTGAGTGCCTCACCA |
| B03 | ACCCTTGAGTATGACA |
| B04 | AGGGTTTCATCGATAC |
| B05 | ATTTCTGGTTGTACGT |
| B06 | CCACGTTTCTCTCGAC |
| B07 | CTAACCCAGTAGCAAC |
| B08 | GAATCGTCATCAGTGT |
| B09 | GCGTGCAGTTGCTAGT |
| B10 | GTCACGGTCTCGACCT |
| B11 | TACTGCCAGTAATTGG |
| B12 | TCGATTTCATAGTCGT |
| C01 | ATAACATGATCGACAC |
| C02 | AAACCCAAGAAACACT |
| C03 | ACGGTCGAGTGGCAGT |
| C04 | AGTAGCTCATTGCAGT |
| C05 | CAAGCTATCACAGCGT |
| C06 | CCCTCTCTCTTGGAAT |
| C07 | CTAGGTAAGTGCATAG |
| C08 | GACGTTACATTAGTCG |
| C09 | GCTTTCGTCAAGTCTG |

| Barcode | Sequence 5' > 3' |
|---|---|
| E01 | CGCCGAGAAACAGACA |
| E02 | AACCTTTAGATGGGCT |
| E03 | ACTTCGCCAAGGTTTC |
| E04 | ATCACAGGTCAACCAT |
| E05 | CACTGTCTCCAGCAGT |
| E06 | CCTCATGAGATCGCTT |
| E07 | CTCCTCCCAAGCGTCC |
| E08 | GAGTGTTGTATGGTAA |
| E09 | GGGACAATCCACAGCG |
| E10 | GTGGGAAAGATACAGC |
| E11 | TCAATTCCAAGACGAC |
| E12 | TGAACGTGTATCGAGG |
| F01 | GATATAACTGGTTGCT |
| F02 | AAGCCATAGCCGAGGT |
| F03 | AGACAGGCACATACCA |
| F04 | ATCGATGGTCGAGATG |
| F05 | CAGCGTGTCCGGACGT |
| F06 | CGAAGGAAGCATTTGT |
| F07 | CTGCAGGCACACTACA |
| F08 | GATCGTAGTCCTCATC |
| F09 | GGGCTCATCCGATAGT |
| F10 | GTTACGAAGCATAGAC |
| F11 | TCAGGGCCACAACGAG |
| F12 | TGAGGAGGTCCCGTGA |
| G01 | GCGGACTGTTTGTCGA |
| G02 | AATCACGAGCTCCACG |
| G03 | AGATCGTCACGTTCGG |
| G04 | ATGACCAGTGACAGGT |
| G05 | CATACTTTCGACCATA |
| G06 | CGATGCGAGCTAAGTG |
| G07 | CTGTAGACACGGATTT |
| G08 | GATTTCTGTCTTGGTA |
| G09 | GGTAATCTCGAACATT |

| | |
|---|---|
| C10 | GTCGCGATCTTCCGGC |
| C11 | TAGGGTTAGTGAACAT |
| C12 | TCGTAGACATGTCAGT |
| D01 | CAGCTAGATGAAGAAG |
| D02 | AACAAAGAGACTGATG |
| D03 | ACTATTCCAAATACGA |
| D04 | AGTTAGCGTAGACAGC |
| D05 | CACATGATCAGTCGAC |
| D06 | CCGTAGGAGACGATCA |
| D07 | CTCAGTCCAAACTAGA |
| D08 | GAGATGGGTACGTGTT |
| D09 | GGAGGTATCAGGAAGC |
| D10 | GTGAGTTAGACCATAG |
| D11 | TATCTTGAGTTTCCAA |
| D12 | TCTCAGCGTACCGCTG |

| | |
|---|---|
| G10 | GTTGCTCAGCGTATAG |
| G11 | TCATGCCCACGAGGAT |
| G12 | TGCACGGGTCTGCGCA |
| H01 | ACAAAGAAGGCAGGTT |
| H02 | AGCGTATCAGATTGGG |
| H03 | ATGGGTTGTGGCGATA |
| H04 | CATGCAATCGGAGGTG |
| H05 | CGGACACAGGATCACG |
| H06 | CGTAGTAAGGGTCTAA |
| H07 | CTTCGGTCAGAGGAAA |
| H08 | GCAGTTAGTGGAACCA |
| H09 | GGTTCTCTCGCTGACG |
| H10 | TAAGCACAGGAGAATG |
| H11 | TCCATCGCAGACCAGA |
| H12 | TGCGGGTGTGCGTCGT |

***Table S5: Nextflex 8bp index barcodes for ID-seq.***

| Barcode | Original sequence | Reversed sequence for primer |
|---|---|---|
| Nextflex 39 | CCTCCTGA | TCAGGAGG |
| Nextflex 40 | CGAACTTA | TAAGTTCG |
| Nextflex 42 | CGCATACA | TGTATGCG |
| Nextflex 43 | CTCAATGA | TCATTGAG |
| Nextflex 46 | GAATCTGA | TCAGATTC |
| Nextflex 47 | CAAGACTA | TAGTCTTG |
| Nextflex 48 | GAGCTGAA | TTCAGCTC |
